# An astrocyte-derived cytokine TIMP-1 restores synaptic plasticity in an Alzheimer’s disease model

**DOI:** 10.1101/2023.03.18.533245

**Authors:** Sukanya Sarkar, Kusumika Gharami, Ramesh Kumar Paidi, B.N. Srikumar, Subhas C. Biswas

**Affiliations:** Cell Biology and Physiology Division, CSIR-Indian Institute of Chemical Biology, 4 Raja S. C. Mullick Road, Kolkata 700 032, India; Department of Neurological Sciences, RUMC, 1735 West Harrison St, Suite Cohn 336, Chicago, IL 60612, USA; Department of Neurophysiology, National Institute of Mental Health and Neuro Sciences (NIMHANS)Hosur Road, Bengaluru – 560029, India

**Keywords:** Astrogliosis, Akt, Amyloid β, cognitive functions, apoptosis, autophagy, TIMP-1, neuroprotection, inflammation, BDNF

## Abstract

The influence of astrocyte-secreted cytokines on neuronal health in Alzheimer’s disease (AD) is poorly understood, despite their increasing recognition as potential molecules for therapeutic targeting. Recently, we demonstrated that an anti-inflammatory cytokine, tissue inhibitor of matrix metalloproteinase-1 (TIMP-1) is robustly secreted by astrocytes early in response to amyloid-β (Aβ) but is reduced in its prolonged presence. Here, we found strikingly diminished levels of TIMP-1 in the brain of six-month-old 5xFAD mice concomitant with high levels of Aβ, versus wild-type mice. Intracerebroventricular injection of TIMP-1 in 5xFAD mice ameliorated their cognitive functions. TIMP-1 not only ensured neuronal viability against apoptosis and aberrant autophagy in the AD model by binding to neuronal CD63 receptors, but also conferred synapse–specific effects. Synaptosomal analysis revealed TIMP-1 elevates dendritic spine size and protein levels, likely by promoting post-synaptic long-term potentiation in hippocampus, independent of pre-synaptic activity. TIMP-1 induced brain-derived neurotrophic factor (BDNF) and BDNF-mediated post-synaptic signaling. Therefore, we identify TIMP-1 as a multifunctional cytokine with distinct protective mechanisms-of-action on neurons and propose it as a promising therapeutic candidate in AD.

**GRAPHICAL ABSTRACT:** 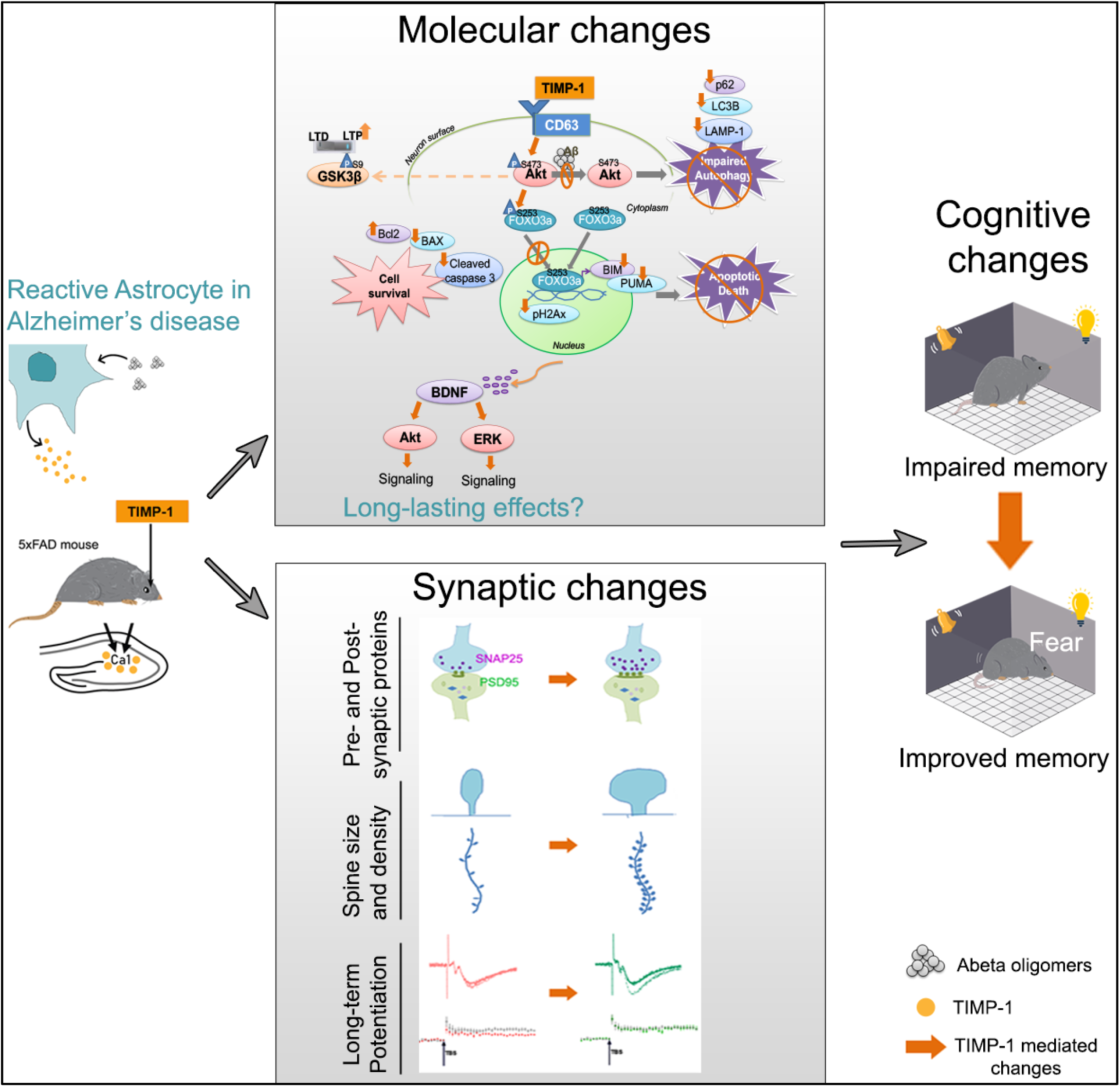

## INTRODUCTION

Alzheimer’s disease (AD) is the most common multi-factorial neurodegenerative disorder (ND) with no effective cure to date. This calls for integrated studies of neurons with glia in AD research. Astrocytes, the most abundant of the glial cells, in addition to their indispensable homeostatic functions in the CNS, are now well-appreciated for being critical in the progress of NDs including AD^1^. Reactive astrogliosis or astrocyte reactivity^2^ in response to any pathological insult to the brain gives rise to a heterogeneous population of reactive astrocyte subtypes with diverse secretomes acting as rich reservoirs of cytokine and chemokine molecules^3, 4^; the exact functions of these inflammatory mediators, especially in a disease scenario, is only partially understood, recently reviewed by us for AD^5^. Exploring the role of individual cytokines and their molecular mechanisms in relation to neuronal health could help explain their implication in progression in pathogenesis, especially in the early stages of NDs and identify novel therapeutic targets.

Advancements in omics studies coupled with single-cell/single nucleus sequencing have unveiled a plethora of reactive astrocyte subtypes existing across AD stages^6–8^. For example, a pathological astrocyte subtype termed ‘disease-associated astrocyte (DAA)’ was detected in the hippocampus of a mouse AD model^6^ while human AD brain post-mortem analysis revealed several pathological astrocyte subtypes with characteristic upregulation in reactivity markers, but downregulation in genes encoding synaptic regulators and glutamate transporters^7, 8^. Moreover, mapping of AD associated astrocyte subtypes onto spatial transcriptomics data from human AD and inflamed mouse brains reveal their regional heterogeneity in brain^9, 10^. However, temporal dynamics of astrocyte reactivity in NDs and related gene expression changes remains poorly explored as recently reviewed^1^. We recently reported the kinetics of Aβ-responsive astrocyte reactivity *in vitro* and interestingly, observed time-dependent differential enrichment of pro- or anti-inflammatory cytokines (secretome) in the astrocyte-conditioned media, among which tissue inhibitor of matrix metalloproteinase 1 (TIMP-1) emerged as one of the major neuroprotective cytokines^11^.

TIMP-1, extensively studied in cancer, has gained some limelight in NDs^12–14^. TIMP-1 is almost exclusively expressed by astrocytes in white matter lesions and plays a neuroprotective role by inhibiting matrix metalloproteinase (MMP) activities^15^. Importantly, a TIMP-1-expressing reactive astrocyte subtype was detected in white matter tracts of a lipopolysaccharide-induced mouse model of neuroinflammation with suggested participation in extracellular matrix remodelling, neuroprotection and remyelination^10^. TIMP-1 and MMP levels prominently differ in the CSF and plasma between ND patients including AD^16–19^ and healthy controls, signifying the importance of studying TIMP-1 in NDs. These reports deal with the role of TIMP-1 in relation to MMP changes. However, several recent reports suggest an emerging MMP-independent role of TIMP-1, reviewed in^20^. TIMP-1 has a two-domain structure – an N-terminal domain responsible for MMP inhibition and a C-terminal domain (CTD) for cytokine-like cellular signaling. The multi-functionality of TIMP-1 arises from the competition among its different binding partners for association with their corresponding attachment domains on TIMP-1, which is influenced by context, the TIMP-1:MMP ratio, and binding affinities of the partners. WhenTIMP-1 levels are higher than MMP levels, free uncomplexed TIMP-1 molecules are available to bind to cognate cell-surface receptors such as CD63^21^, CD82^22^ and LRP-1^23^, through its CTD, inducing downstream signaling pathways. Furthermore, during pathological insults, when the levels of MMPs are higher than those of TIMP-1, MMP-9 activity increases^24^; more pro-MMP9s, initially occupying TIMP-1’s CTD, are cleaved into MMP-9 and therefore, the CTD becomes free for binding and signaling through CD63 via integrinβ1, MAPK^21^, FAK^25^ and PI3K/Akt^26, 27^ pathways. Thus, TIMP-1 under both these situations induces intracellular signaling, few of which are implicated in TIMP-1-mediated beneficial effects against CNS insult^25, 27^. Therefore, from our previous work and those of others, TIMP-1 emerges as an astrocyte-derived neuroprotective molecule, but its main mechanisms of action on neurons in AD from an intracellular signaling perspective remain incompletely understood.

In this study, we explored a plausible multifunctional role of TIMP-1 in an advanced transgenic model of AD – the 5xFAD mice. We observed alleviation of cognitive deficits in 5xFAD mice which drastically lacks endogenous TIMP-1, with TIMP-1 treatment. We detected the underpinning intracellular mechanisms, especially the synaptic changes driven by TIMP-1 in rescuing cognitive malfunctions in 5xFAD mice; identifying the major cellular mediators of its modus operandi and providing evidence for its possible therapeutic implication in AD.

## RESULTS

### TIMP-1 injection rescues cognitive deficits in 5xFAD mice

5xFAD mice start developing Aβ plaques at 2 months^28^ with extensive reactive astrogliosis, microgliosis by 4 months, and eventually synaptic and neuronal loss^29^. We first checked the levels of TIMP-1 (secreted form, 25 kDa^30^) in 5xFAD mice versus wild type C57/BL6 mice (WT) across different age groups. TIMP-1 was markedly reduced in hippocampal (Figures 1A and 1B, Figures S1A and 1B) and cortical tissues (Figures S1C and S1D) of 2-month (M) and 4M - 10M-old 5xFAD mice compared to age-matched WT groups as determined by western blot (WB) analyses. There was a four-fold reduction in TIMP-1 expression compared to WT from 6M onwards in the hippocampus of 5xFAD mice. Because of the substantial loss of TIMP1 in AD mice, we next examined whether its replacement with exogenously supplied protein would ameliorate their behavioral deficits. A total amount of 5 ng of TIMP-1 dissolved in artificial cerebro-spinal fluid (aCSF) was injected intracerebroventricularly (i.c.v.) in each mouse; the dose used is based on our previous work^11^ and comparable to the TIMP-1 concentration in human CSF^16^ (see materials and methods). Injection and all subsequent experiments were performed in 6M-old WT and 5xFAD mice. 5xFAD mice injected with aCSF were used as the vehicle control. Since the level of endogenous TIMP-1 was dramatically reduced in 5xFAD mice, we predict that ‘injected’ TIMP-1 (Figure 1C) directly contributes to the changes that are subsequently observed, without a significant contribution from endogenous TIMP-1. Diffusion of TIMP-1 by 14 days following i.c.v. injection, in the hippocampal and cortical regions of 5xFAD mice was confirmed by immunofluorescence studies (Figures 1D and 1E).

**Figure 1:**
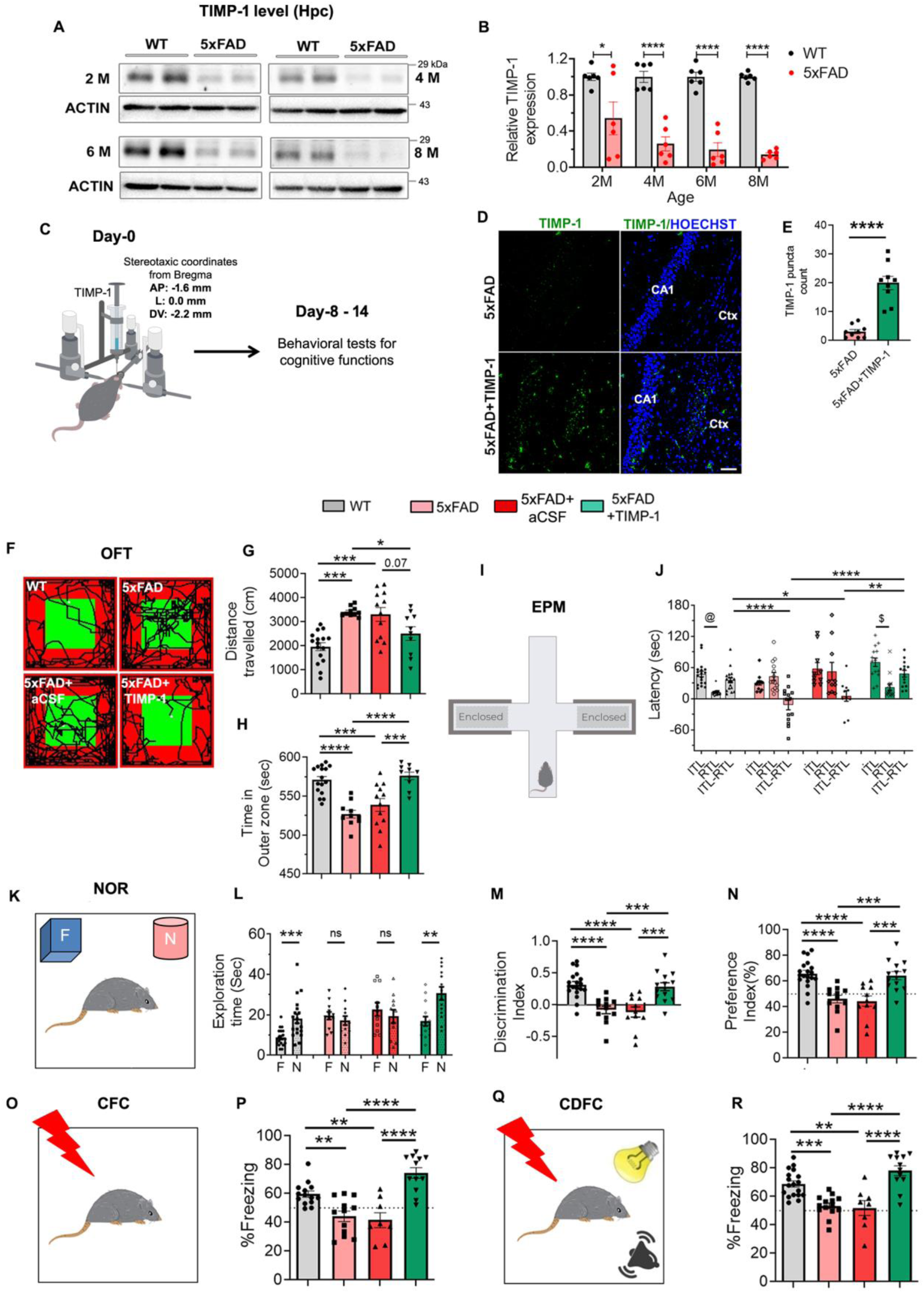
Exogenous TIMP-1 injection corrects cognitive deficits in 5xFAD mice. **A**, Representative immunoblots showing endogenous TIMP-1 levels in wild type (WT) and 5xFAD mice in the hippocampus at 2 months (M), 4M, 6M and 8M of age (also refer Figures S1A and S1B). **B**, Quantification of TIMP-1 levels were normalized to actin. Relative expression was calculated w.r.t. WT in each age group (N=6 mice/group; two-tailed unpaired t-test, mean±s.e.m.). **C**, Schematic of stereotactic injection of TIMP-1 (5 ng) and behavioral experiment schedule. **D**, Representative immunofluorescence images of coronal sections showing hippocampal regions from 5xFAD and 5xFAD+TIMP-1 mouse brains stained for TIMP-1 (green) and nuclei (blue), 14 days post TIMP-1 injection; scale bar-50 µm. **E**, Quantification for TIMP-1 immunoreactivity in the hippocampus shown as TIMP-1 puncta count (N=3 mice/group, analyzed 3 slices/mouse brain; two-tailed unpaired t-test, mean±s.e.m.). **F-R**, Battery of behavioral tests performed with four groups of mice – WT, 5xFAD, 5xFAD+aCSF and 5xFAD+TIMP-1 starting on Day-8 following intracerebroventricular TIMP-1 injection to assess cognitive functions. **F**, Representative infra-red images of the path travelled by a mouse in each group in the Open-field test (OFT). **G**, Average total distance travelled (cm) in the whole open arena and **H**, Average time spent (sec) in the outer zone of the arena in 10 min of OFT. One-way ANOVA, Tukey’s post hoc, mean±s.e.m. **I**, Experimental set-up for elevated-plus maze (EPM) test. **J**, Average latency (sec) of mouse to the closed arm of EPM recorded on Day-1 as initial transfer latency (ITL) and on Day-2 as retention transfer latency (RTL) and their difference (ITL-RTL). For ITL versus RTL within each group, two-tailed unpaired t-test, mean±s.e.m; @P<0.0001, #P<0.001. For comparison of ITL-RTL among groups, One-way ANOVA, Tukey’s post hoc, mean±s.e.m. **K**, Experimental set-up for Day-2 of Novel object recognition (NOR) test with one familiar (*F*) and one novel (*N*) object for **L-N**.**L**, Average exploration time (sec) with *F* versus *N* within 5 min on Day-2 of NOR; two-tailed unpaired t-test, mean±s.e.m. **M**, Discrimination index and **N**, Preference index (%) calculated among groups from Day-2 data of NOR, One-way ANOVA, Tukey’s post hoc, mean±s.e.m. For OFT, EPM and NOR also see Figure S2. **O**, Experimental set-up for Day-1 of contextual fear conditioning (CFC) test. **P**, Percentage freezing of mice on Day-2 of CFC. **Q**, Experimental set up for Day-1 of Cue-dependent fear conditioning (CDFC) test and **R**, Percentage freezing of mice on Day-2 of CDFC, One-way ANOVA, Tukey’s post hoc, mean±s.e.m. for **P** and **R**. *P<0.05, **P<0.01, ***P<0.001 and ****P<0.0001; ns, not significant. Each point of the Scatter plots with bar represents one animal. Number of animals and exact P values in each test are given in Table S1.

Seven days following TIMP-1 injection into 5xFAD mice, a battery of behavioral tests was performed to determine the effect of TIMP-1 on cognitive behaviors. In an open-field test (OFT), both 5xFAD and artificial cerebro-spinal fluid (aCSF)-injected 5xFAD mice displayed hyperlocomotion (Figures 1F and 1G; FigureS2A) and decreased thigmotactic behavior (Figures 1F and 1H and Figure S2B), suggesting increased anxiety compared to WT mice, that was significantly attenuated by TIMP-1 (5 ng) as represented by the average distance travelled (in cm) and the time spent in the outer zone (sec) by mice in each group in the 10 min period of testing. In the elevated plus maze (EPM) test (schema in Figure 1I), there was no significant difference between the initial transfer latency (ITL) on Day-1 and retention transfer latency (RTL) on Day-2 in the 5xFAD and 5xFAD+aCSF groups while in both WT and TIMP-1-treated 5xFAD groups RTLs were reduced significantly compared to ITLs (Figure1J, Figure S2C). Mean ITL-RTL values are additionally represented for each group (Figure 1J; Figure S2D), showing that TIMP-1 treatment significantly improved ITL-RTL scores in 5xFAD mice (positive value) compared to the untreated 5xFAD group (negative value). These observations indicate that 5xFAD mice display better memory retention following TIMP-1 treatment. In the novel object recognition (NOR) test (schema for Day-2 with novel (N) and familiar (F) objects in Figure1K), both 5xFAD and 5xFAD+aCSF mice spent significantly a shorter time with novel objects than that with familiar objects on Day-2 (Figure 1L; Figure S2E), further evidenced by negative values for the Discrimination index (DI; Figure 1M) and <50% score in the Preference index (PI; Figure 1N). TIMP-1 treatment significantly improved the novel object exploration time of 5xFAD mice on Day-2 (Figure 1L; Figure S2E), indicated further with positive DI values (Figure 1M) and >50% PI scores (Figure 1N). TIMP-1 therefore ameliorated long-term recognition memory in 5xFAD mice. Figures 1O and 1Q show schemas for contextual fear conditioning (CFC) and cue-dependent fear conditioning (CDFC) (Day-1 acquisition) tests respectively. In both the tests, the average %freezing of a mouse in 8 min encompassing the four test sessions on Day-2 (see ‘Materials & Methods’) was calculated. On Day-2 of CFC, the 5xFAD and 5xFAD+aCSF mice displayed a diminished retention of context-dependent fear memory, that is, reduced %freezing value versus WT mice. This was significantly improved in the TIMP-1-treated 5xFAD group (Figure 1P). Similarly, cue-dependent fear memory was disrupted in the 5xFAD and 5xFAD+aCSF mice on Day-2 of CDFC and corrected by TIMP-1 injection (Figure 1Q). The results indicate a role of TIMP-1 in better retention of fear memory that involves appropriate coordination between hippocampus and amygdala. Additionally, we assessed the changes in Aβ plaque deposition 14 days post TIMP-1 treatment. Both Aβ plaque burden and plaque number were significantly reduced by TIMP-1 injection in 5xFAD mice (Figures S3A-S3C). Altogether, these data suggest that TIMP-1 is detected at significantly reduced levels in 6M-old 5xFAD mice and coincides with enhanced Aβ plaque deposition and cognitive deficits, both of which are reversed by recombinant TIMP-1 injection.

### TIMP-1 binds to CD63 on neurons mediating Akt phosphorylation

The next question that arises is ‘how’ TIMP-1 exerts its beneficial effects in 5xFAD mice. To answer that, we first explored the ability of TIMP-1 to bind to a neuronal surface receptor in order to initiate any neuroprotective signaling mechanism. CD63 is a cell-surface receptor (a member of the tetraspanin family) responsible for TIMP-1-mediated Akt signaling^31^ (Schema in Figure 2A). We have shown that Akt is involved in TIMP-1-induced neuroprotection against Aβ toxicity^32^. Immunofluorescence staining for TIMP-1 (red) and CD63 (green) 16 h following TIMP-1 (100 ng/ml) addition to primary cortical neurons showed significantly enhanced co-localization between the two proteins (Figure 2B) on the neuronal surface quantified by Pearson correlation coefficient (Figure 2C).

**Figure 2:**
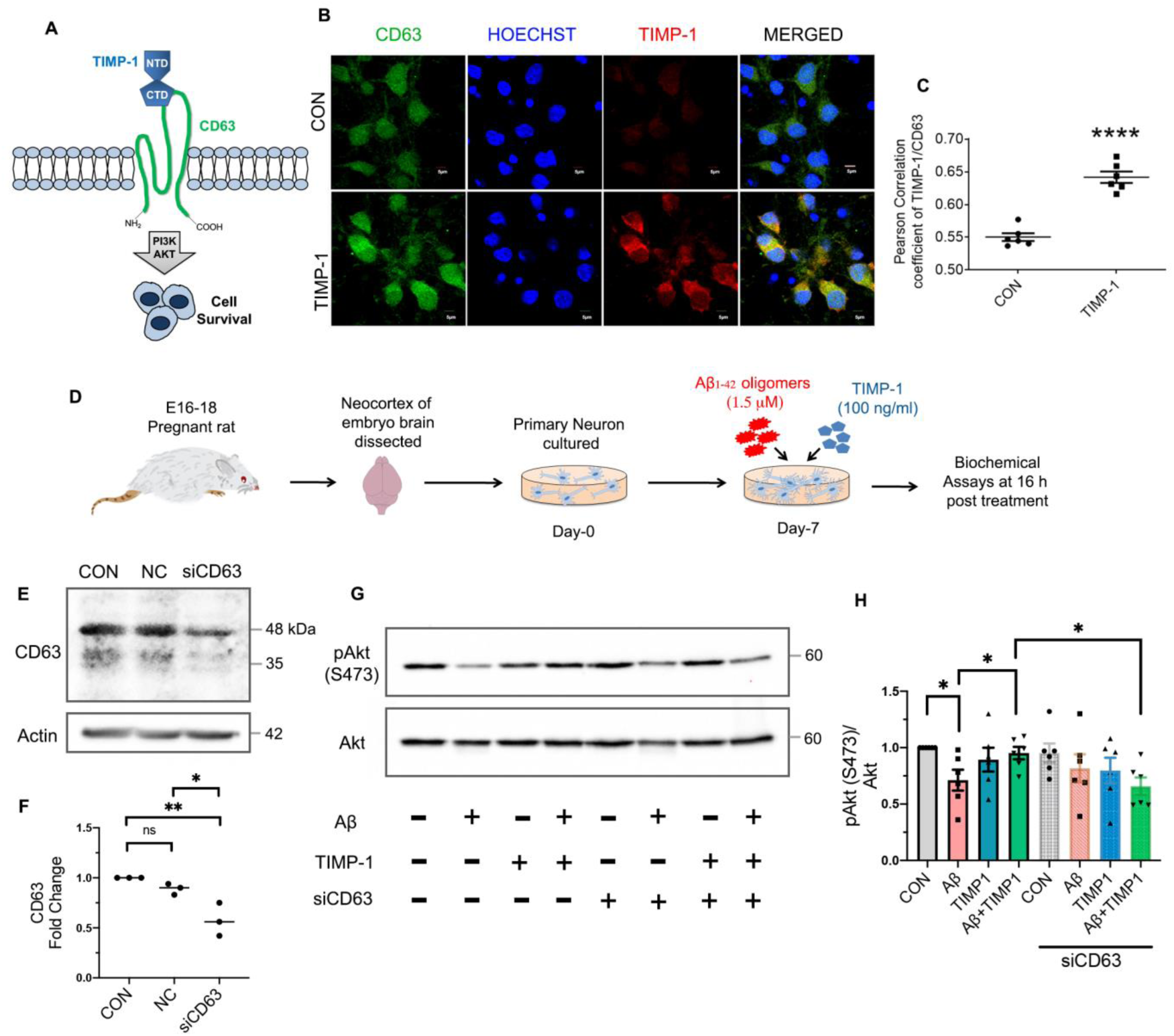
TIMP-1 binds to CD63 on neurons and enhances Akt phosphorylation. **A,** Schema showing TIMP-1 binding to CD63 receptor activates PI3K/Akt pathway. **B**, Representative immunofluorescence images for CD63 (green), Hoechst (blue) and TIMP-1 (red) in primary cortical neurons 16h post-treatment with or without (CON) TIMP-1 (100 ng/ml). Merged panel shows colocalization (yellow) between CD63 and TIMP-1. Scale bar, 5 µm. **C**, Colocalization analysis by calculating Pearson correlation coefficient between TIMP-1 and CD63 (N = 3 independent experiments, 2 wells/experiment; two-tailed unpaired t-test, mean±s.e.m; ****P<0.0001). **D**, Schema for primary neuron culture and treatment. **E**, Representative immunoblot for CD63 from primary cortical neurons treated with or without siCD63 for siRNA validation along with neurons treated with Negative control siRNA (NC). **F**, Densitometric analysis and calculation of CD63 fold change in NC and siCD63-treated neurons w.r.t. CON utilizing actin for loading control normalization (N=3 independent experiments; One-way ANOVA, Tukey’s post hoc, mean±s.e.m; *P<0.0209, **P<0.0051). **G**, Representative immunoblots for pAkt(S473) and corresponding Akt from primary neurons, with or without CD63 downregulation, treated with or without Aβ (1.5 µM), TIMP-1 or co-treated with Aβ+TIMP-1. **H**, Densitometric analysis and calculation of fold change of pAkt(S473)/Akt w.r.t. CON for N=6 independent experiments; two-tailed unpaired t-test, mean±s.e.m. *P=0.0109 CON vs. Aβ, *P=0.0499 Aβ vs. Aβ+TIMP-1 and *P=0.0114 Aβ+TIMP-1(no siCD63) vs. Aβ+TIMP-1 (with siCD63).

To investigate the role of CD63 in TIMP-1-mediated Akt phosphorylation in primary neurons (schema in Figure 2D), we acquired CD63 siRNA (siCD63) and validated its efficacy (Figures 2E and 2F) 72 h following treatment. Expectedly, Akt phosphorylation was attenuated at 16 h post Aβ treatment compared to control cells and was reversed significantly by TIMP-1 addition to Aβ- treated neurons (Figures 2G and 2H). Interestingly, upon CD63 downregulation, the improvement in Akt phosphorylation induced by TIMP-1 in Aβ-treated neurons was lost. Hence, we interpret that TIMP-1 binds to CD63 receptor on neuronal surface which is required for TIMP-1-mediated Akt phosphorylation at Ser473.

### TIMP-1 restricts nuclear translocation of FOXO3a and related apoptotic signaling induced by Aβ

Aβ treatment of cortical neurons causes dephosphorylation of Akt and consequently of its downstream target FOXO3a. Dephosphorylated FOXO3a(S253) translocates to the nucleus increasing transcription of pro-apoptotic genes *Bim*^33^ and *Puma*^34^ (Schema in Figure 3A). We found that TIMP-1 restored the phosphorylation of FOXO3a(S253) in Aβ-treated primary neurons at 16 h following treatment (Figures 3B and 3C) while the total FOXO3a and actin levels remained unaltered across the treatment conditions. Consistently, FOXO3a puncta (green) were increased in the nuclei (blue) of Aβ-treated neurons16 h post-treatment, indicating that FOXO3a’s nuclear localization was attenuated by TIMP-1 treatment (Figures 3D and 3E). FOXO3a-induced pro apoptotic proteins BIM (Figures 3F and 3G) and PUMA (Figures3F and 3H) were significantly upregulated in Aβ-treated neurons, but this was markedly reduced under Aβ+TIMP-1 co-treatment (Figures 3F-H). Interestingly, anti-apoptotic protein Bcl-2 levels decreased in Aβ-treated neurons, and elevated upon TIMP-1 treatment (Figures 3F and 3I), while another anti-apoptotic protein, Bcl-xL, remained unchanged across the treatment conditions (Figures 3F and 3J). Notably, Bcl-2 and Bcl-xL levels are differentially regulated in response to TIMP-1 in different disease models and stages^12, 35, 36^. Furthermore, pro-apoptotic BAX (Figures 3F and 3K), DNA damage marker pH2Ax (Figures 3F and 3L) and apoptotic effector cleaved Caspase3 (Figures 3F and 3M) levels were down-regulated in Aβ+TIMP-1-treated cells in contrast to Aβ-treated neurons. Thus, TIMP-1 protects neurons against Aβ toxicity by inhibiting apoptosis induced by the Akt/FOXO3a/BIM/PUMA axis.

**Figure 3:**
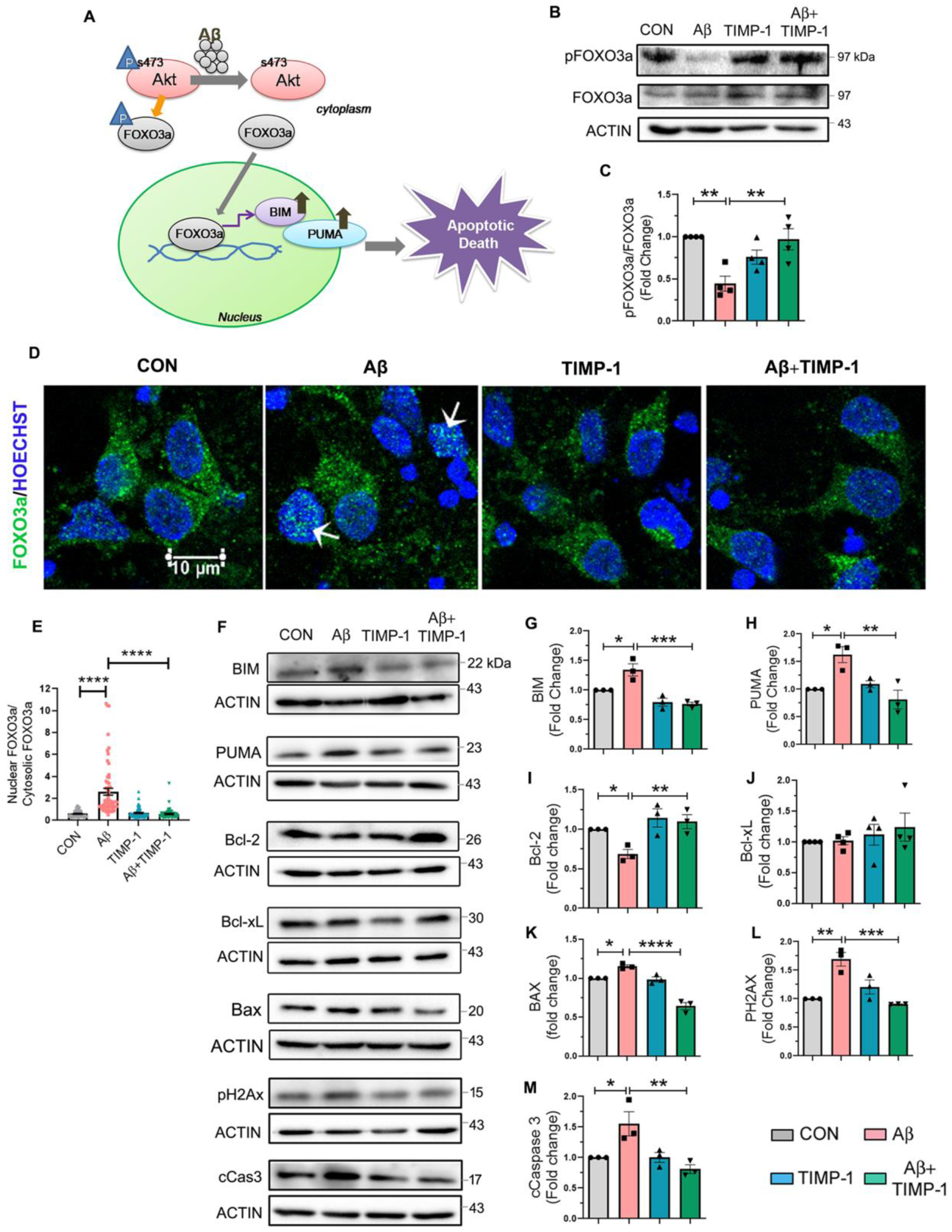
TIMP-1 protects primary cortical neurons against apoptosis. **A**, Schema showing that Aβ treatment of neurons causes Akt dephosphorylation at S473 and downstream nuclear translocation of FOXO3a, upregulating apoptosis as previously reported^30, 31^. **B**, Representative immunoblots for pFOXO3a(S253) and corresponding total FOXO3a and Actin at 16 h post treatment with or without TIMP-1 (100 ng/ml) in presence or absence of Aβ(1.5 µM) in primary cortical neurons. **C**, Densitometric analysis followed by calculation of fold change for pFOXO3a/FOXO3a w.r.t. control (CON) (N = 4 independent experiments; One-way ANOVA, Tukey’s post hoc, mean±s.e.m.). **D**, Representative merged immunofluorescence images for FOXO3a (green) and nuclei stained with Hoechst (blue). White arrows indicate green dots for FOXO3a in the nuclei in the Aβ-treated neurons. **E**, Quantification of nuclear translocation of FOXO3a by ImageJ, N = 3 independent experiments, 20 cells/experiment; One-way ANOVA, Tukey’s post hoc, mean±s.e.m. **F**, Representative panel of immunoblots from primary neuron lysates probed with pro and anti-apoptotic proteins as indicated along with the corresponding loading control or actin from the same blot. **G-M**, Densitometric analyses followed by calculation of fold change w.r.t. CON for BIM (**G**), PUMA (**H**), Bcl-2 (**I**), Bcl-xL (**J**), BAX (**K**), pH2Ax (**L**) and cCas3 (**M**), (N = 3 independent experiments; One-way ANOVA, Tukey’s post hoc, mean±s.e.m.).*P<0.05, **P<0.01, ***P<0.001, ****P<0.0001. cCas3, cleaved Caspase 3. Also see Figure S4.

Next, in order to study the effect of TIMP-1 on apoptosis *in vivo*, we performed TUNEL assays in coronal brain sections that revealed a significant increment in the number of TUNEL-positive (apoptotic) cells in the hippocampal regions of 5xFAD mice compared to WT mice. This was significantly reduced 14 days post TIMP-1 treatment (FiguresS4A and S4B). Furthermore, TIMP-1 abrogated the high levels of pH2Ax in 5xFAD mice as determined by WB analyses of whole hippocampal tissue (Figures S4C and S4D) and immunofluorescence studies of hippocampal (Figures S4E and S4F) and cortical sections (Figures S4G and S4H). Together, these results corroborate our *in vitro* findings.

### TIMP-1 corrects impaired autophagic flux in Aβ-treated primary neurons and in 5xFAD mice

Our lab has shown earlier that Aβ treatment of primary cortical neurons simultaneously induces both apoptosis and aberrant autophagy showing accumulation of autophagic proteins in neurons, and leading to impaired autophagic flux^37, 38^ (schema in Figure 4A). Therefore, here we investigated the effect of TIMP-1 on impaired autophagic flux induced by Aβ in neurons. We found increased levels of ATG5 (an upstream marker for elongation of autophagosomes, Figures 4B and 4C), p62 (cargo adaptor and the most crucial determinant of autophagic flux, Figures 4D and 4E) and LC3B-II (Figures 4F and 4G) by WB analyses of primary neuron lysates following Aβ treatment. Lysosome numbers were up-regulated, identified by lysosome-marker LAMP1 levels in Aβ-treated neurons, indicating that although autophagy was induced by Aβ, the flux was eventually impaired (Figures 4H and 4I). TIMP-1 addition to Aβ-treated neurons improved autophagic flux by reversing the levels of ATG5, p62, LC3BII and LAMP1 to control levels (Figures 4B-4I). Immunofluorescence analyses of p62 (Figures 4J and 4K), LC3B (Figures 4L and 4M) and LAMP1 (Figures 4N and 4O) levels under the same conditions 16h post-treatment corroborated the WB results. Hence, both apoptosis and autophagy are regulated by TIMP-1 possibly through phosphorylation of a common upstream kinase, Akt, protecting cells against Aβ toxicity.

**Figure 4:**
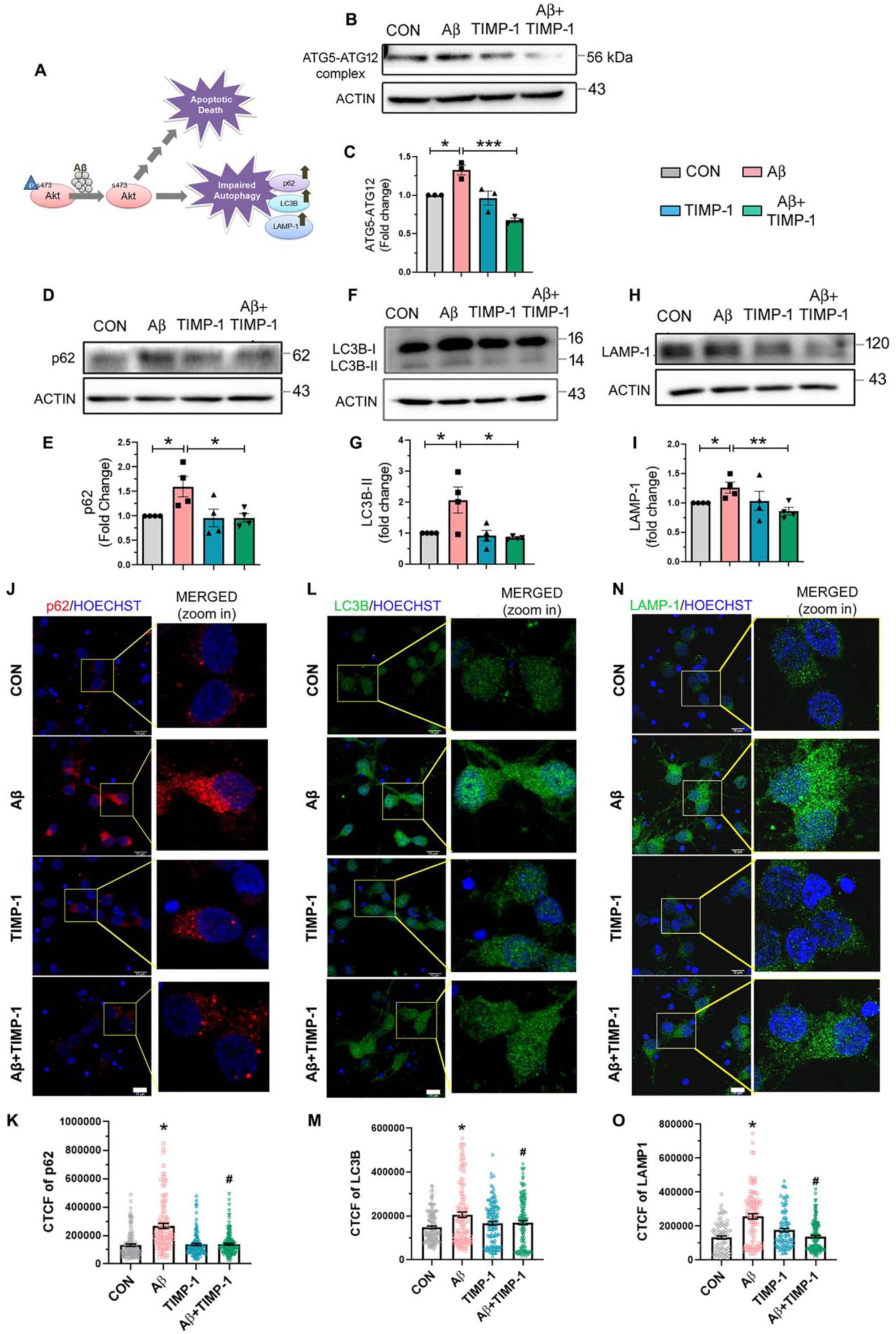
TIMP-1 addition ameliorates autophagy flux in Aβ-treated neurons. **A**, Schematic showing Aβ-induced Akt inactivation causes impaired autophagy flux, in addition to apoptosis induction, in neurons^34^. **B**, Representative immunoblot for ATG5-ATG12 complex along with corresponding actin from primary neurons at 16 h post treatment with or without TIMP-1 (100 ng/ml) in presence or absence of Aβ (1.5 µM). **C**, Densitometric analysis and calculation of fold change in ATG5-ATG12, normalized to actin, w.r.t. control cells (CON), (N = 3 independent experiments; One-way ANOVA, Tukey’s post hoc, mean±s.e.m.). **D-I**, Representative immunoblots and corresponding fold changes w.r.t. CON forp62 (**D-E**), LC3B-II (**F-G**) and LAMP1 (**H-I**). N = 4 independent experiments; One-way ANOVA, Tukey’s post hoc, mean±s.e.m.*p<0.05, **p<0.01, ***p<0.001 for **B-I**.**J,L,N**, Representative merged immunofluorescence images for p62 (**J**, red), LC3B (**L**, green) and LAMP1 (**N**, green) in primary neurons treated similarly as in western blot, at 16 h posttreatment; nuclei visualized with Hoechst (blue); scale bar, 10 µm. A zoomed-in panel is shown for each protein to highlight the change in fluorescence intensities within individual cells. Corrected total cell fluorescence (CTCF) calculated for p62 (**K**), LC3B (**M**) and LAMP1 (**O**) from N = 3 independent experiments, analyzed 90-150 cells/treatment condition; One-way ANOVA, Tukey’s post hoc, mean±s.e.m; *P<0.0001 versus CON, ^#^P<0.0001 versus Aβ. Also see Figure S5.

*In vivo*, TIMP-1 upregulated inhibitory phosphorylation of ULK-1 at Ser757 that likely prevented autophagy induction in 5xFAD mouse hippocampus (Figures S5A and S5C) and cortex14 days post treatment (Figures S5B and S5D). Elevated levels of p62 (Figures S5E-S5H), LC3B (Figures S5I-S5L) and LAMP1 (Figures S5M-S5P) were detected in 5xFAD mouse brain by immunofluorescence studies, indicating that although autophagy was induced in 5xFAD mice, the pathway was eventually impaired, leading to increased accumulation of autophagosomes and lysosomes that was ameliorated by TIMP-1 treatment. These *in vivo* findings thus replicate the *in vitro* results.

### TIMP-1 enhances pre and post-synaptic protein expression in 5xFAD mice

Synapse loss has long been considered the strongest correlate of cognitive impairment in AD^39^. Therefore, we investigated the synaptic underpinnings of the cognitive improvement we observed in TIMP-1-treated 5xFAD mice. Pre and post-synaptic protein levels, common determinants of synaptic health, were assessed in whole tissue and in synaptosomes (Figure 5A). TIMP-1 injection in 5xFAD mice induced a two-fold increase in pre-synaptic SNAP25 fluorescence intensities in CA1 hippocampi (Figures 5B and 5C) and cortices (Figures S6A and S6B) compared to untreated 5xFAD mice, 14 days post-treatment. Similarly, post-synaptic marker PSD95 intensities were also enhanced in CA1 hippocampi (Figures 5D and 5E) and cortices (Figure S6C and S6D) of 5xFAD+TIMP-1 mice. WB analyses of whole hippocampal tissue from TIMP-1-treated 5xFAD mice further corroborated the increase in SNAP25 (Figures 5F and 5G) and PSD95 (Figures 5H and 5I) levels compared to 5xFAD group. However, no significant changes were observed in these synaptic protein levels in whole cortices among the these animal groups (Figures S6E-H). No significant change was observed across groups for additional synaptic markers in hippocampus Synaptophysin-1 (pre-synaptic) and Homer-1 (post-synaptic) (Figures S6I and S6J).

**Figure 5:**
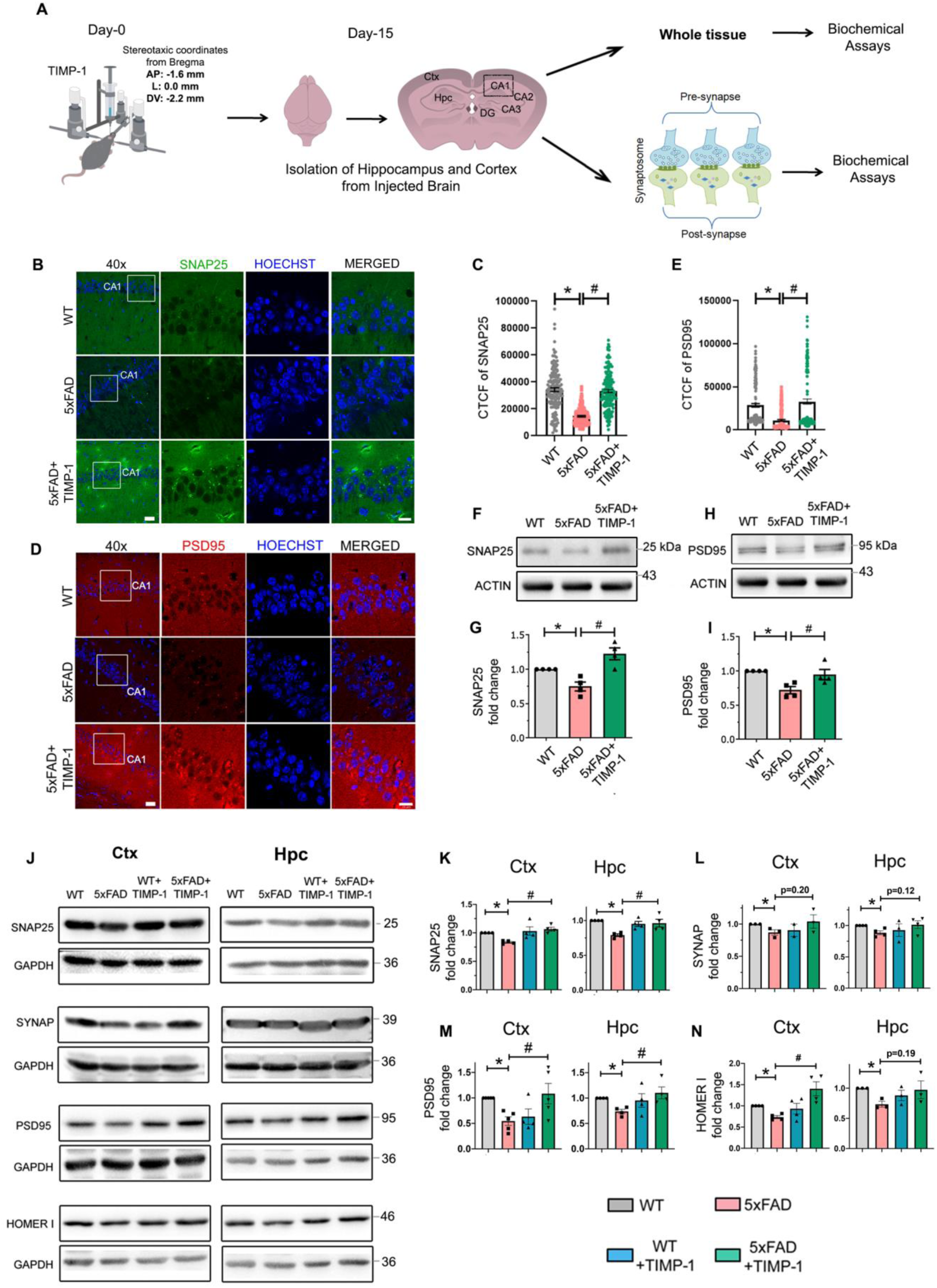
TIMP-1 enhances synaptic protein expression in 5xFAD mice. **A,** Schematic showing methodology for whole tissue and synaptosome isolation from hippocampus (Hpc) and cortex (Ctx) of TIMP-1-injected mouse brain. **B, D,** Representative immunofluorescence images for SNAP25 (green, **B**) and PSD95 (red, **D**) in coronal sections of WT, 5xFAD and 5xFAD+TIMP-1 hippocampus; scale bar, 30 µm at 40x magnification (white box indicates the region selected for further zoom-in). Zoomed-in merged images are shown with nuclei stained with Hoechst (blue); scale bar, 15 µm. CTCF calculated for SNAP25 (**C**) and PSD95 (**E**), N= 3 animals/group, n = 50 cells/animal from Hpc quantified from 40x images; One-way ANOVA, Tukey’s post hoc, mean±s.e.m. *P<0.0001, #P<0.0001 for **C** and **E**. Refer to Figures S6A-S6D for cortical expressions of the above two proteins. Representative immunoblots for SNAP25 (**F**) and PSD95 (**H**) from Hpc whole tissue lysates collected 14 days post TIMP-1 injection. Densitometric analyses and fold changes for normalized SNAP25 (**G**) and PSD95 (**I**) w.r.t. to WT group. N = 4 animals/group; One-way ANOVA, Tukey’s post hoc, mean±s.e.m. **G**, *P=0.0449, ^#^P=0.0011;**I**, *P=0.012, ^#^P=0.0351. See Figures S6E-S6H for WB results for these two proteins from whole cortex tissue. Also see Figures S6I-S6J for HOMER-1 and Synaptophysin-1 (SYNAP) expressions in whole Hpc. **J**, Representative immunoblots for synaptic proteins from synaptosomes isolated separately from Ctx and Hpc. Densitometric analyses and fold changes w.r.t. WT were calculated for pre-synaptic markers SNAP25 (**K**, Ctx *P=0.0012, ^#^P=0.0001; Hpc, *P=0.0049, ^#^P=0.0154) and SYNAP (**L**, Ctx, *P=0.0354; Hpc, *P=0.0103), and post-synaptic markers PSD95 (**M**, Ctx *P=0.0007, ^#^P=0.0397; Hpc, *P=0.0008, ^#^P=0.0260) and HOMER-1 (**N**, Ctx *P=0.0003, ^#^P=0.0074; Hpc, *P=0.0053). N = 3-5 independent synaptosome isolations/group, 6-7 pooled animals/synaptosome isolation/group; mean±s.e.m. Statistical tests used for **K-N** are given in Table S1. **F** and **H** show representative images of proteins from the same blot and hence same actin. Similarly in **J**, PSD95 (Hpc) and HOMER1 (Hpc) are from same blot, hence same GAPDH.

In order to delineate the changes mediated by TIMP-1 specifically at the synapse, we utilized synaptosomal preparations. Pre-synaptic nerve terminals in synaptosomes often remain attached to the post-synaptic membrane and post-synaptic density proteins^40^. WB analyses with viable synaptosomes from the four treatment groups of mice – WT, 5xFAD, WT+TIMP-1 and 5xFAD+TIMP-1 revealed that the expressions of pre-synaptic (Figures 5J-5L) and post-synaptic (Figures 5J, 5M and 5N) markers were significantly lower in 5xFAD mice compared to WT mice both in the cortical and hippocampal regions. Fourteen days post TIMP-1 treatment, the level of SNAP25 (Figures 5J and 5K) and PSD95 (Figures 5J and 5M) were significantly elevated in cortices and hippocampi of 5xFAD mice. TIMP-1 did not significantly increase Synaptophysin-1 levels in cortical or in hippocampal 5xFAD synapses (Figures 5J and 5L). Unlike the whole tissue results, Homer-1 levels were significantly higher in the 5xFAD+TIMP-1 mouse cortex but not in hippocampus (Figures 5J and 5N). No significant changes were observed in synaptic protein expressions with TIMP-1 treatment of WT mice. Altogether, these results suggest that TIMP-1 was able to increase specific synaptic protein levels in 5xFAD mice.

### TIMP-1 ameliorates deficits in synaptic spine density and spine size in 5xFAD mice

Dendritic spines, possessing the majority of excitatory synapses, are highly dynamic in their size, shape and density, and these changes are strongly correlated with learning and memory functions^41^. Golgi-cox staining and imaging of brain slices from 5xFAD mice (Figure 6A), 14 days post TIMP-1 injection, revealed a significant increase in spine density (number of spines/10 µm) in hippocampal (CA1) (Figures 6B and 6C) and cortical regions (Fig 6B and 6D) compared to untreated 5xFAD mice. The spine density remained unaltered between the WT and WT+TIMP-1 groups.

**Figure 6:**
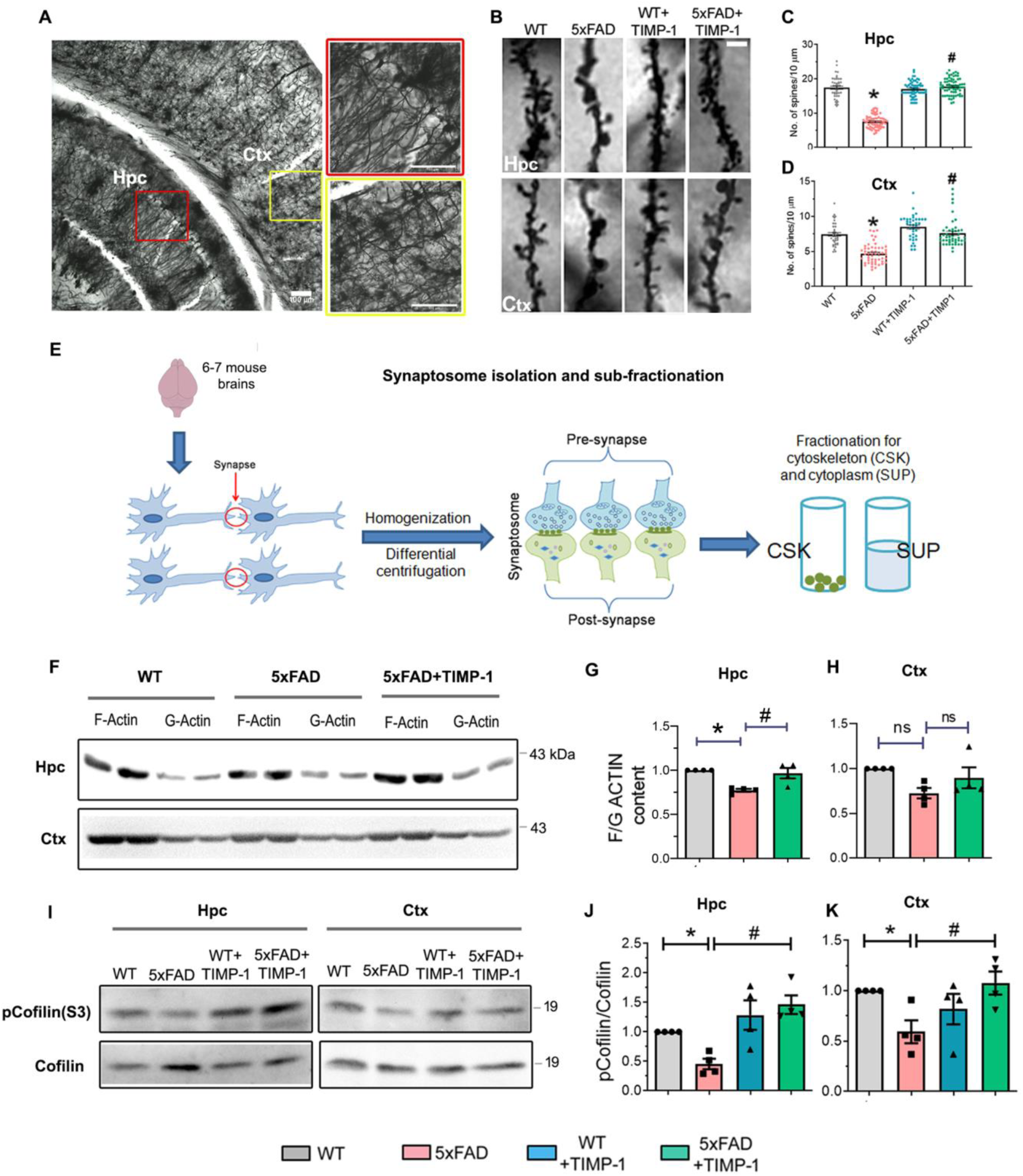
TIMP-1 improves spine density and size in the 5xFAD mouse. **A**, Representative coronal section of a mouse brain showing the CA1 region of hippocampus (Hpc, red box) and the cortex (Ctx, yellow box) targeted for spine density calculation following Golgi-cox staining (scale bar-100 µm). **B**, Representative images of dendritic segment showing spines in Hpc (upper panel) and Ctx (lower panel) from the four treatment groups, 14 days post TIMP-1 injection; scale bar, 2 µm. **C-D**, Quantification of the number of spines/10 µm from Hpc (**C**, *P<0.0001 versus WT; ^#^P<0.0001 versus 5xFAD) and Ctx (**D**, *P<0.0001 versus WT, ^#^P<0.0001 versus 5xFAD). N = 3 animals/group, 40-80 dendritic stretches/group for each brain region; One-way ANOVA, Tukey’s post hoc, mean±s.e.m. **E**, Schema for synaptosome isolation followed by fractionation for cytoskeleton (CSK) and cytoplasmic/supernatant (SUP) fractions. **F**, Representative immunoblots for filamentous (F, in CSK fraction) actin and globular (G, in SUP fraction) actin from Hpc and Ctx synaptosomes. **G-H**, Densitometric analysis and fold-change for F/G actin content w.r.t. WT from Hpc (**G**, *P=0.0038, ^#^P=0.0101) and Ctx (**H**, ns=not significant). N=4 independent synaptosome isolations/group, 6-7 pooled animals/synaptosome isolation/group; One-way ANOVA, Tukey’s post hoc, mean±s.e.m. No change in F/G actin ratio was observed between WT and WT+TIMP-1 groups (refer Figure S7). **I**, Representative immunoblots for pCofilin(S3) and Cofilin from Hpc and Ctx synaptosomes. **J-K**, Densitometric analysis and fold change for pCofilin/Cofilin from Hpc (**J**,*P=0.0125, ^#^P=0.0002) and Ctx (**K**,*P=0.0313, ^#^P=0.0127). N=4 independent synaptosome isolations/group, 6-7 pooled animals/synaptosome isolation/group; One-way ANOVA, Tukey’s post hoc, mean±s.e.m.

Besides spine density, spine shrinkage is a well-studied abnormality in AD^42^. Changes in spine size are regulated by dynamic alterations in the cytoskeletal actin content^43^ quantifiable by the ratio of filamentous(F) versus globular(G) forms of actin in synaptosomes. We fractionated viable synaptosomes into the cytoskeletal fraction containing F actin and the supernatant fraction containing G actin (Figure 6E). The F/G actin ratio was significantly reduced in hippocampal synaptosomal sub-fractions from 5xFAD mice compared to the WT group. TIMP-1 treatment elevated the spine cytoskeletal actin content in 5xFAD mice, indicated by an increased F/G actin ratio (Figures 6F and 6G) but did not affect the spine actin content in WT mice (Figure S7). Furthermore, no significant alteration was observed in the cortical synaptosomal actin contents among the treatment groups (Figures 6F and 6H). Spine size and maturation are correlated with the stability of the actin cytoskeleton. Phosphorylation of Cofilin (an actin-binding protein) is a major regulator for stabilization of the actin cytoskeleton and thereby promotes spinal maturation followed by spine enlargement^44^. Phosphorylated Cofilin(S3) levels were approximately 50% lower in 5xFAD mice compared to WT mice, both in the hippocampus (Figures 6I and 6J) and cortex (Figures 6I and 6K). This was significantly increased by TIMP-1, indicating that TIMP-1 helps in inducing spine maturation and consequent enhancement in spine size, more specifically in the hippocampus.

### TIMP-1 induces BDNF and related signaling in 5xFAD mice

TIMP-1 improved synaptic properties in 5xFAD mice. Hence, we questioned whether this effect was direct or indirect by induction of a neurotrophin, namely brain-derived neurotrophic factor (BDNF), an important regulator of spine properties^45^ and synaptic plasticity^46^. We found a four fold reduction in levels of Pro-BDNF, the precursor of mature BDNF, in whole hippocampal lysates from 5xFAD mice compared to WT animals. Fourteen days post TIMP-1 treatment, Pro BDNF levels increased in 5xFAD mice to reach WT levels (Figures 7A and 7B). In contrast, TrkB, the receptor for BDNF, both in its full length (fl-TrkB) (Figures 7C and 7D) and truncated (t-TrkB) (Figures 7C and 7E) isoforms were not significantly altered in the hippocampus. CREB is a transcription factor that promotes *Bdnf* gene expression and the pCREB(S133)/CREB/BDNF axis is known to be dysregulated in AD^47^. In consonance with the BDNF results, we observed a similar increase in pCREB(S133)/CREB expressions in 5xFAD+TIMP-1 mice versus 5xFAD mice (Figures S8A and S8B). This led us to investigate the effect of TIMP-1 on BDNF-TrkB signaling pathways implicated in synaptic plasticity, namely, the PI3K/Akt and ERK/MAPK pathways^48^. Akt phosphorylation, which is stimulated by BDNF^49^, was elevated at Ser473 in TIMP-1-injected 5xFAD mice (Figures 7F and 7G) compared to untreated 5xFAD mice, which is consistence with our *in vitro* results, and additionally at Thr308 (Figures 7H and 7I). Phosphorylation levels of ERK1 (Figures 7J and 7K) and ERK2 (Figures 7J and 7L), were unchanged indicating that the ERK pathway remained unaltered.

**Figure 7.**
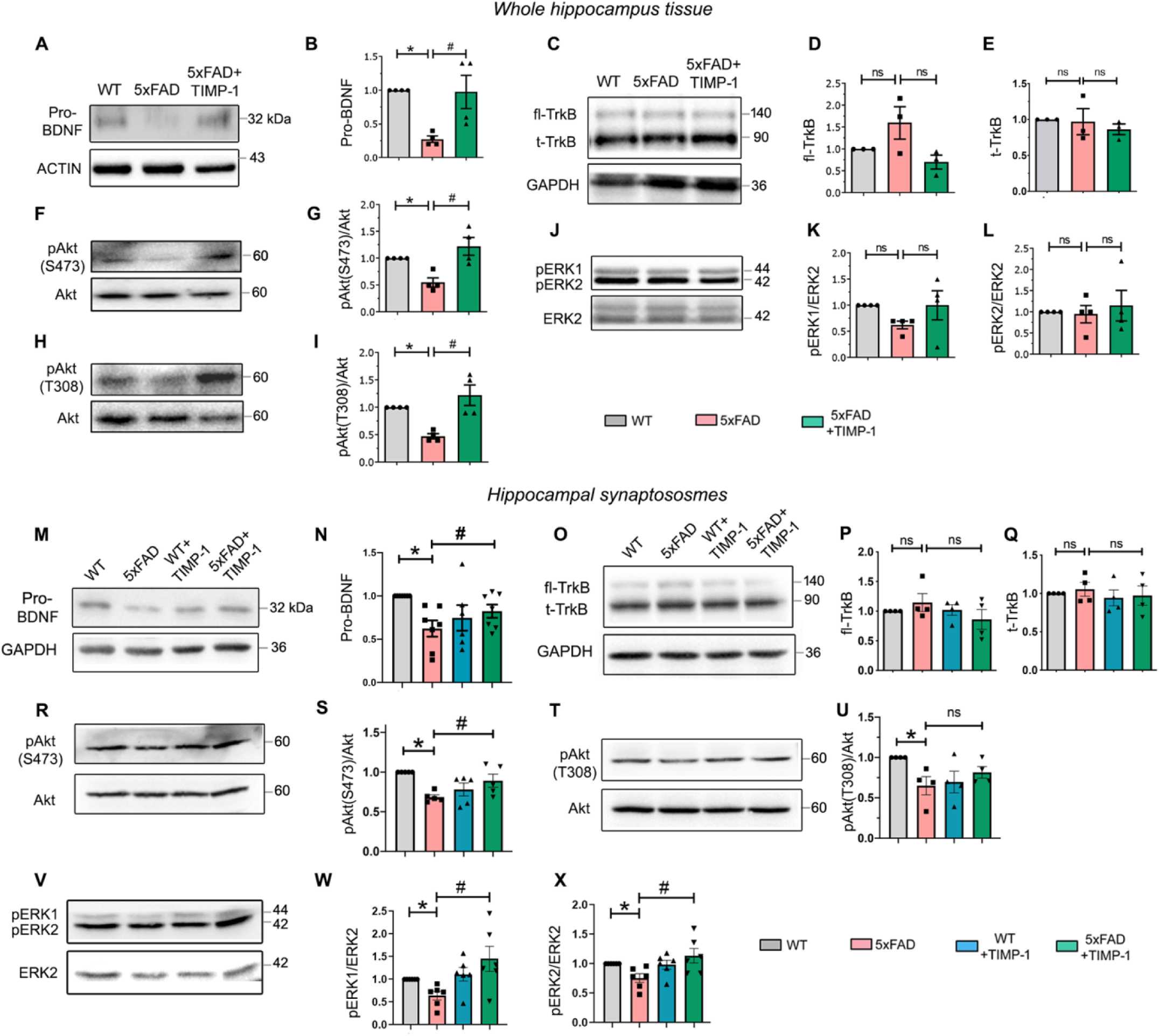
BDNF and related signaling pathways are induced by TIMP-1 treatment of 5xFAD mice. **A-L**, Western blot (WB) data from whole hippocampal (Hpc) tissue lysates 14 days post TIMP-1 treatment. Representative immunoblots for Pro-BDNF (**A**), TrkB (**C**), pAkt(S473) (**F**), pAkt(T308) (**H**) and pERK1/2 (**J**) with corresponding loading controls. Densitometric analyses and normalized fold-change w.r.t. WT for Pro-BDNF (**B**), fl-TrkB (**D**), t-TrkB (**E**), pAkt(S473)/Akt (**G**), pAkt(T308)/Akt (**I**), pERK1/ERK2 (**K**) and pERK2/ERK2 (**L**). N = 3-4 mice/group; One-way ANOVA, Tukey’s post hoc, mean±s.e.m. **M-X,** WB data from Hpc synaptosomes 14 days post TIMP-1 treatment. Representative blots for Pro-BDNF (**M**), TrkB (**O**), pAkt(S473) (**R**), pAkt(T308) (**T**) and pERK1/2 (**V**) with corresponding loading controls. Densitometric analyses and normalized fold-change w.r.t. WT for Pro-BDNF (**N**),fl-TrkB (**P**), t-TrkB (**Q**), pAkt(S473)/Akt (**S**), pAkt(T308)/Akt (**U**), pERK1/ERK2 (**W**) and pERK2/ERK2 (**X**). N=7 independent synaptosome isolations/group, Mixed effect analysis, Tukey’s post hoc for **N**; N=4-6 independent synaptosome isolations/group, One-way ANOVA, Tukey’s post hoc for **P,Q,S,U,W,X**. Each synaptosome isolation (N) pooled from 6-7 animals for each group. All data expressed as mean±s.e.m. fl-TrkB, Full-length TrkB; t-TrkB, truncated TrkB. Also see Figure S9 for WB results from cortical synaptosomes. *P<0.05 versus WT, ^#^P<0.05 versus 5xFAD, ns=not significant. Exact P values for this figure are given in Table S1. R and V are obtained from the same blot.

Pro-BDNF level in isolated synaptosomes from adult mice is a measure of the release-ready pre synaptic secretory (vesicular) pool of BDNF’s precursor form^50^. We detected a reduction of almost 50% in pro-BDNF levels at hippocampal (Figures 7M and 7N) and 20% at cortical (Figures S9A and S9B) synapses in 5xFAD mice. TIMP-1 elevated the levels significantly in both regions, 14 days post-treatment. However, TrkB (fl-TrkB and t-TrkB) remained unaltered across treatment groups in hippocampal (Figures 7O-7Q) and cortical (Figures S9C-S9E) synaptosomes, as in total tissues. Further, we found increased phosphorylation of Akt at Ser473 (Figures 7R and 7S) but not at Thr308 (Figures 7T and 7U) in hippocampal synaptosomes from TIMP-1-treated 5xFAD mice compared to untreated 5xFAD animals, indicating that although Akt signaling is induced at synapses by TIMP-1 treatment, not all of Akt’s active sites are equally phosphorylated. In cortical synapses, we detected a similar increase in levels of pAkt(S473) expression in the 5xFAD+TIMP-1 group compared to the 5xFAD group (Figures S9F and S9G), but there was no difference in the pAkt(T308) levels between groups (Figures S9H and S9I). Conversely, unlike total hippocampal levels, we observed significant reductions in pERK1/ERK2 (Figures 7V and 7W) and pERK2/ERK2 (Figures 7V and 7X) levels in hippocampal synapses from 5xFAD mice that were elevated upon TIMP-1 treatment, indicating that alterations in ERK activation could be detected at the synaptic level, but not at the whole tissue level. A similar recovery in both pERK1/ERK2 (Figures S9J and S9K) and pERK2/ERK2 (Figures S9J and S9L) levels was observed at cortical synapses after TIMP-1 treatment.

### TIMP-1 enhances basal synaptic transmission and long-term potentiation in the hippocampal Schaffer collateral-CA1 synapses of 5xFAD mice

BDNF induces long-term potentiation (LTP) in hippocampus^51^. LTP/LTD regulation forms the basis of synaptic plasticity that underlies learning and memory^52^. Hence, we investigated whether TIMP-1 treatment affected LTP induction in the 5xFAD model. A schema of the *ex vivo* electrophysiology at Schaffer collateral-CA1 synapses in mouse hippocampus is shown in Figure 8A. First, we assessed whether basal synaptic transmission was affected at Schaffer collateral-CA1 synapses in 5xFAD mice compared to WT mice by measuring the input-output (I-O) relationship. Figure 8B shows representative fEPSP traces at increasing current stimulus intensities (20 – 300 µA with 20 µA increments), from each experimental group. In 5xFAD slices, the fEPSP response at all current intensities was lower than that in WT slices. TIMP-1 enhanced the fEPSP slope values in 5xFAD mouse significantly at several current intensities, the highest being at 260 µA (Figure 8C). Area under curve (AUC) analysis for Figure 8C corroborated the reduced basal transmission observed in 5xFAD mice and its significant enhancement upon TIMP-1 treatment (Figure 8D). Next, we studied the effect of TIMP-1 treatment of 5xFAD mice on short-term plasticity by determining the paired-pulse facilitation (PPF) ratio at Schaffer collateral-CA1 synapses. PPF helps in evaluating the alterations in pre-synaptic activity and neurotransmitter release probability by delivering two consecutive stimulating pulses with a short inter-stimulus interval (ISI). The PPF ratio remained unaltered among the experimental groups (Figure 8E), further corroborated by AUC calculation (Figure 8F). Thus, the probability of pre-synaptic neurotransmitter release did not differ between WT and 5xFAD mice at hippocampal SC-CA1 synapses, with no further change upon TIMP-1 treatment.

**Figure 8:**
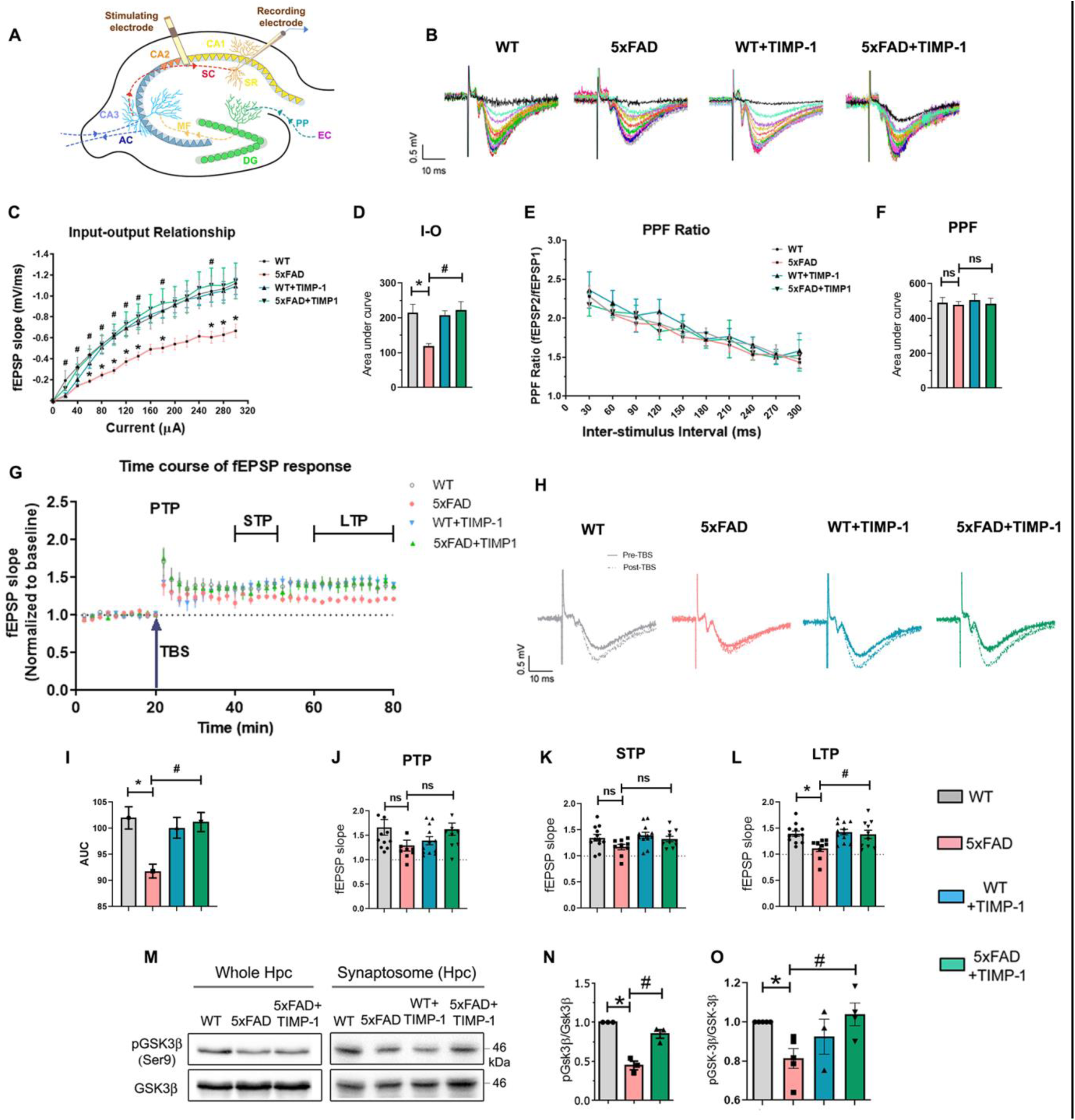
Deregulated basal synaptic transmission and long-term potentiation at Schaffer collateral-CA1 synapses in 5xFAD mice are enhanced by TIMP-1 through inhibition of hippocampal GSK3β activity, but the paired pulse facilitation ratio remains unaffected in all the groups. **A**, Schema showing regions of the hippocampus – CA1, CA2, CA3 and Dentate gyrus (DG) and the Entorhinal cortex (EC). The dotted lines mark the connecting fibres – Schaffer collateral (SC), Mossy fibres (MF), Perforant pathway (PP), Associational commissural fibres (AC). Stratum radiatum (SR) is where the recording electrode is placed in the CA1 region. **B**, Representative field excitatory post-synaptic potentiation (fEPSP) traces of slices from WT, 5xFAD, WT+TIMP and 5xFAD+TIMP-1 treatment groups with increasing current stimulus strength (20 – 300 µA with 20 µA increments). **C**, Input-output (I-O) relationship (fEPSP slopes along a step-wise rise in stimulus intensity) representative of the mean fEPSP value of all the slices in each group at each stimulus current strength from 20 – 300 µA (n=9-15 slices/group, N=5-6 mice/group, Two-way ANOVA, mean±s.e.m.; *P<0.05, ^#^P<0.05. **D**, Area under curve (AUC) from **C**, One-way ANOVA, Tukey’s post hoc, mean±s.e.m. *P=0.0153, ^#^P=0.0103. **E**, Paired pulse facilitation (PPF) ratio along increasing inter-stimulus interval (30 to 300 ms with 30 ms increments), Two-way ANOVA, mean±s.e.m. **F**, AUC from **E**, n=9-15 slices/group, N=5-6 mice/group, One-way ANOVA, Tukey’s post hoc, mean±s.e.m. **G**, Time course of fEPSP responses with 20 min pre-TBS and 60 min post-TBS. Note the time segments post-TBS demarcated for calculating post-tetanic potentiation (PTP), short-term potentiation (STP) and long-term potentiation (LTP). **H**, Representative fEPSP tracings of slices from the four groups. Solid line represents pre-TBS response (baseline) and dotted line represents post-TBS response. Note the decrease in the magnitude of LTP in 5xFAD (pink) compared to WT (grey) and increase in LTP in the 5xFAD+TIMP-1 (green) group compared to 5xFAD. **I**, AUC from **G,** One-way ANOVA, Tukey’s post hoc, mean±s.e.m. *P= 0.0088, ^#^P= 0.0207. **J-L**, Bar graphs for PTP (3min post-TBS) (**J**), STP (21^st^-30^th^ min) post-TBS (**K**), LTP (41^st^-60^th^ min post-TBS) (**L**); n=9-12 slices/group, N=5-6 mice/group, One-way ANOVA, Tukey’s post hoc, mean±s.e.m. *P=0.0165, ^#^P=0.0355. **M**, Representative immunoblots for pGSK3β(S9) from whole tissue and synaptosomal lysates from hippocampus (Hpc), 14 days following TIMP-1 injection. **N-O**, Densitometric analysis and fold change w.r.t. WT for pGSK3β/GSK3β from whole tissue (**N**,*P=0.0003, ^#^P=0.0017) and synaptosome (**O**, *P=0.0202, ^#^P=0.0094). N=3mice/group for **N**, N=4 independent synaptosomal isolations for **O**; One-way ANOVA, Tukey’s post hoc, mean±s.e.m. *P versus WT, ^#^P versus 5xFAD for **C-D,I,L,N-O**. ns=not significant.

We evaluated synaptic potentiation at three different plasticity time points – Post-tetanic potentiation (PTP, 3min post theta burst stimulation (TBS)), short-term potentiation (STP, 21^st^- 30^th^ min post TBS) and LTP (41^st^-60^th^ min post TBS i.e. early-phase LTP). Figure 8G shows the time course of normalized fEPSP responses of the treatment groups 20 min before and 60 min after TBS. Figure 8H shows representative tracings of pre and post-TBS fEPSPs in each group. AUC of the total time course of fEPSP responses in 5xFAD slices was significantly decreased compared to WT slices, while AUC in the 5xFAD+TIMP-1 group was comparable to WT and WT+TIMP-1 groups (Figure 8I). LTP induction was found to be significantly lower in the 5xFAD hippocampal slices compared to WT mice, in consistence with previous studies^29, 53^. Importantly, TIMP-1 injection in 5xFAD mice significantly enhanced the level of LTP induction while LTP induction remained similar between WT and WT+TIMP-1 slices (Figure 8L). No significant change was detected in PTP (Figure 8J) or STP (Figure 8K) responses among the treatment groups.

GSK3β is an important negative regulator of LTP implicated in synaptic changes in the brain^54^ and plays a critical role in AD^55^. Since we found enhanced LTP induction with TIMP-1 treatment of 5xFAD mice, we hypothesized that TIMP-1 may have a role in GSK3β signaling. We found increased GSK3β phosphorylation at Ser9, indicating GSK3β inhibition, in whole tissue (Figures 8M and 8N) and synaptosomal lysates from hippocampi of TIMP-1-treated 5xFAD mice compared to low inhibitory phosphorylation levels in untreated 5xFAD mice, indicating GSK3β activation (Figures 8M and 8O).

## DISCUSSION

Our research^32^, in consonance with recent reports^5, 56, 57^, indicates that reactive astrocytes in AD can be an excellent source of disease-modulating cytokines whose detailed mechanism-of-actions are incompletely understood. In this study, we revealed the underlying key molecular mechanisms of TIMP-1, an astroglia-derived, Aβ-responsive anti-inflammatory cytokine, identified by us earlier^32^, to have effects on neuroprotection, synaptic plasticity and cognitive recovery in AD. We were intrigued to find that endogenous TIMP-1 levels are strikingly diminished in the hippocampus of a transgenic model of AD, the 5xFAD mouse, compared to WT mice across age. Interestingly, intra-cerebroventricular injection of exogenous TIMP-1 in this mouse model not only improved their long-term hippocampus-dependent contextual memory (explicit memory), but also rescued fear memories (implicit memory). While TIMP-1 ensured neuronal viability by regulating two prominent, yet intertwined, cell-death pathways – apoptosis and autophagy by binding to its cognate receptor CD63, it also improved synaptic integrity and function – the direct cellular correlates of cognitive recovery. The latter effect may be attributed to an enhanced expression of BDNF and BDNF-associated signaling as well as to GSK3β inhibition, especially at synapses, following TIMP-1 injection to 5xFAD mice.

In 5xFAD mice, we detected lower endogenous TIMP-1 levels than in WT mice across age, the earliest being at 2 months, coinciding with an inherent early Aβ accumulation in this model^28^. One study showed that TIMP-1 increases significantly at 4 months in 5xFAD mice but falls to the WT level at 6 months of age^58^. Interestingly, in the CSF of AD patients TIMP-1 levels were found to be lower in comparison to healthy individuals^18, 19^. A strong body of evidence suggests a biphasic regulation of TIMP-1 in response to acute versus chronic inflammatory insults associated with HIV-associated neurocognitive disorders (HAND)^13^. In the study, TIMP-1 was regarded as a neuroprotective molecule that decreases in CSF and tissue samples from chronic HAND patients compared to healthy individuals but increases when an acute inflammatory insult is given to primary astrocytes in culture^12, 14^. The latter is consistent with our report on acute treatment of astrocytes with Aβ^32^. Possibly, the early secretion of TIMP-1 in response to Aβ is an attempt by the astrocytes to protect and repair neuronal damage due to the insult but in case of a prolonged presence of Aβ, as in aging 5xFAD mice, TIMP-1-mediated neuroprotection is lost due to the downregulation of TIMP-1 by an unknown mechanism. Interestingly, transcriptomes of neurons derived from human induced pluripotent stem cells generated from skin fibroblasts of AD patients carrying a mutation in the *PSEN1* gene show downregulation of *Timp-1* gene^59^, emphasizing TIMP-1’s neuroprotective significance and alluding to a possible connection with AD-associated genes.

Our previous work shows that TIMP-1 can activate Akt^32^ but the downstream cellular pathway/s effective against AD remained unexplored. Here, we demonstrate that CD63 is the specific receptor responsible for TIMP-1-driven Akt activation in Aβ-treated primary neurons utilizing co localization and RNAi techniques. TIMP-1 via CD63 induces Akt signaling in many different cell types^26, 27, 31^. CD63 is probably the most important binding partner for intracellular TIMP-1 signaling as opposed to other competitive interactors, especially when TIMP-1 is in excess and unbound^24^ like the exogenous addition in our study. While we do not completely disregard a possible effect of TIMP-1 injection on MMP activities in 5xFAD mice or a subsequent contribution towards the beneficial effects mediated by TIMP-1, we predict that the MMP independent role of TIMP-1, as a signaling molecule, majorly contributes to the long-term effects of TIMP-1 in our model system *in vivo* (discussed below). Since AD is a proteopathic disease where autophagy and apoptosis cooperate in mediating neurotoxicity^37, 38^ and Akt is a common upstream regulatory kinase for both apoptosis and autophagy^60^, TIMP-1 likely mediates neuroprotection by stimulating this common regulatory point (phosphorylating Akt) for the interconnected cell death pathways in our AD models. Nevertheless, regulation of both apoptosis and autophagy has direct implications in cognitive functioning which have not been explored here.

Progressive changes in synaptic structure and function often precede the actual neuronal loss and memory deficits in AD^61–63^. We find that TIMP-1 not only ensures overall neuronal viability, but also protects synaptic structures. It improves pre and post-synaptic protein levels as well as spine density and size in 5xFAD mice compared to untreated ones. The actin cytoskeleton plays a pivotal role in spine dynamics, both from structural and functional perspectives. Notably, loss of F-actin occurs selectively at synapses at very early ages in a mouse model of AD (APPswe/PS1ΔE9), well ahead of the onset of pathological hallmarks and memory deficits in this model^64^. We show that the F/G actin ratio was significantly increased in hippocampal synapses with TIMP-1 treatment in 5xFAD mice as opposed to untreated 5xFAD animals. We further provide evidence that TIMP-1 reduces the hyperactivated levels of cofilin (dephosphorylated Cofilin) in isolated synaptosomes from 5xFAD mice. Hyperactive cofilin in the 5xFAD mouse is capable of forming rod-shaped actin/cofilin bundles in the cytoplasm of dendritic spines, blocking axonal trafficking and is responsible for synaptic dysfunctions leading towards synaptic loss in AD^65–68^. These events are likely prevented by TIMP-1. Our observations suggest that TIMP-1 has an important role in regulating synaptic dynamics. However, not all the synaptic markers are sensitive to TIMP-1 treatment in 5xFAD mice which is consistent with previous reports on differential responses of synaptic proteins in AD brain^69–71^. Moreover, we did not explore whether the final synaptic recovery promoted by TIMP-1 directly prevented neuronal death or whether there are two independent mechanisms at play. Nevertheless, our work draws attention towards the beneficial roles of an astrocyte-secreted factor, TIMP-1 on synaptic structure and function in neurodegenerative diseases, beyond the body of evidence of several critical physiological effects of astrocyte-secreted proteins on synapses^72–77^.

Reduced release of BDNF in brain and the circulation are key determinants of pathological conditions in brain including AD^78, 79^. In consistence with these reports, we detected significantly diminished levels of BDNF and BDNF-mediated signaling at hippocampal synapses in 5xFAD mice. TIMP-1 not only strongly elevates BDNF levels in 5xFAD mice, but also enhances BDNF/TrkB-directed downstream Akt and ERK signaling pathways at these synapses. Our observations strengthen the ameliorative role of TIMP-1 specifically at synapses as demonstrated by its ability to induce the expression of a well-recognized synaptically crucial neurotrophin.

A key finding of our study is the ability of TIMP-1 to enhance LTP in the 5xFAD mouse hippocampus. Previous studies have mostly shown cytokine-driven LTP suppression^80, 81^, constituting cytokine-driven neuroinflammation, as a contributing event in AD; here we demonstrate a cytokine-mediated beneficial upregulation of LTP in an AD model. Interestingly, TIMP-1 *albeit* via extracellular MMP-9 inhibition, is required for both LTP maintenance^30^ and late-phase LTP inhibition^82^, the latter seen as a strategy to impede non-target late-phase LTP induction and to ensure consolidation of ongoing plasticity. We show that TIMP-1 contributes to early-phase LTP induction as a cell-signaling molecule triggering intracellular pathways. However, whether TIMP-1 has a local effect at synapses as a part of astrocyte-neuron communication is an open question. Interestingly, both PPR and PTP that measure changes in short-term plasticity and the probability of neurotransmitter release from pre-synaptic terminals^83^ remained unchanged across the treatment groups. Therefore, TIMP-1-driven enhancement of LTP in the hippocampus of the 5xFAD mouse model is probably independent of pre-synaptic activity.

GSK3β is associated with PSD, regulates NMDAR-dependent synaptic plasticity^84^ and NMDAR trafficking^85, 86^. Increased GSK3β activity in AD^87^ has the potential to negatively impact synaptic plasticity, in turn learning and memory^88^. GSK-3α and β are present within synapses^88^; more precisely, an electron microscopic study shows GSK-3β labelling of postsynaptic densities at dendritic spines^89^. Extrapolating from electrophysiological data, we predict that our observation of TIMP-1-mediated GSK3β inhibition in 5xFAD mice likely occurs at the post-synaptic compartment and not at the pre-synaptic terminal.

Our data demonstrate an ameliorative role of TIMP-1 at behavioral, viability and synaptic levels. However, the results collectively point to hippocampus as the main seat of action. Notably, in some instances we did not find significant change in cortical whole tissues or synaptosomes. These results may be evidence of differential regulation in different brain regions or due to utilization of heterogeneous whole cortex tissues comprising affected and unaffected synaptic regions for WB analyses. In *in vitro* experiments, TIMP-1 alone does not seem to have any significant effect on the levels of apoptotic or autophagic proteins in control neuronal cells at 16 h following treatment. This may be because the primary neurons are maintained in Neurobasal supplemented with B27 media where growth factors (such as insulin) are present that maximizes the levels of proteins (such as pAkt, pFOXO3a etc.) and the addition of TIMP-1 cannot alter it further. However, Aβ treatment, at the dose used, alters the levels of these proteins (such as reducing levels of pAkt, pFOXO3a etc.) probably by compromising the beneficial effects of growth factors from media. Hence, we get an observable change in protein expressions in Aβ-treated cells with TIMP-1 possibly because TIMP-1 can maintain a sustained beneficial effect (discussed below). We also found a null effect of TIMP-1 injection in WT mice in our *in vivo* studies; one explanation may be the presence of an optimum level of TIMP-1 in adult WT mice contrary to 5xFAD mice, hence exogenous TIMP-1 cannot further induce any observable change in WT mice.

We show that TIMP-1 acts via Akt and BDNF signaling and the longest time we observe these effects is up to two weeks from the time of TIMP-1 injection *in vivo*. The question that comes to mind is how TIMP-1 regulates such long-term effects, especially the synaptic changes. Interestingly, a recent report demonstrates that BDNF can modulate its own expression in cortical neurons by inducing a transcriptional positive feedback loop. It further shows that CREB family transcription factors along with coactivator CBP/p300 are the major regulators of *Bdnf* gene expression following an initial BDNF/TrkB signaling giving rise to a BDNF/TrkB autoregulatory loop^90^. Considering our findings, we may speculate that TIMP-1-induced BDNF expression and the subsequent BDNF/TrkB signaling in 5xFAD mice help in maintaining a sustained expression of BDNF through an autonomous pathway (independent of TIMP-1) involving CREB activation, which possibly contributes to the long-term effects of TIMP-1. Another intriguing point is that the *Timp-1* gene is nested within *Synapsin* gene; this nesting is highly conserved across vertebrates and invertebrates separated by millions of years of evolution^91^. The evolutionarily conserved nesting of *Timp-1* gene within *Synapsin* is of particular significance since Synapsin protein is an important synaptic regulator^92^ and there are prior reports demonstrating co-regulation of genes nested with *Timps*^93^. Moreover, TIMP-1 KO studies demonstrate that TIMP-1 is essential in maintaining synaptic plasticity^94^ and its sequestration prevented spine maturation^30^. Interestingly, LTP induction also causes an upregulation in TIMP-1 mRNA^95^ and protein levels^30^. One paper also shows a sustained expression of TIMP-1 up to 14 days in astrocytes (consistent with our effect duration with TIMP-1 injection) where they suggest that the long-lasting expression of TIMP-1 post kainate treatment (a rat model of seizure) may be due to TIMP-1’s ability as a trophic factor to induce a cascade of interactions between neurons and non-neuronal cells, where trophic factors can regulate each other’s expressions^96^. In our study, since we observe enhanced LTP induction with TIMP-1 treatment in 5xFAD mice, one possibility is that this LTP may in turn give rise to enhanced *Timp-1* gene expression and the subsequent sustained trophic effects contributing towards long-term changes. Together, we speculate that one of these two mechanisms – BDNF autoregulation independent of TIMP-1 and TIMP-1 autoregulation or both may be responsible for the long-lasting effect of TIMP-1 mediated changes in 5xFAD mice which can have important clinical implications. It would be further interesting to address the kinetics of Akt phosphorylation driven by TIMP-1 to divulge whether it is directly mediated by TIMP-1’s signaling through CD63, is a downstream effect of TIMP-1-directed BDNF/TrkB signaling, or whether both occur simultaneously.

In conclusion, TIMP-1 emerges as a multifunctional cytokine released by reactive astrocytes at early stages in AD which not only protects neurons against apoptosis and autophagic deregulation, but has a direct implication on several regulators of synaptic plasticity. This explains the mechanistic basis of TIMP-1-mediated cognitive recovery in 5xFAD mice. Three major revelations are (1) the notable extent of BDNF induction observed upon exogenous TIMP-1 injection to 6-month-old 5xFAD mice that have greatly reduced levels of endogenous TIMP-1, (2) the prominent induction of TIMP-1-driven LTP in 5xFAD mice and (3) the action of TIMP-1 as a regulatory hub for multiple interconnected pathways impacting neuronal properties. This presents a new perspective of TIMP-1’s role in the CNS, yet consistent with the evidence of its multifunctionality as recently reviewed^20, 24^. These findings broadly indicate that a specific astrocyte-secreted cytokine can impact cell functions at different levels which may be seen as a biological strategy^20, 97^ undertaken by astrocytes to ensure flexibility to adapt to changes in homeostasis in NDs including AD.

AD affects one-third of the population above 65 years of age and is the most debilitating form of dementia^98^. Numerous failures in anti-AD therapy make it imperative to investigate new pathophysiological mechanisms or molecules in the early stages of AD and utilizing astrocyte secreted anti-inflammatory factors promises an exciting approach. Our work raises the possibility of utilizing astrocyte-derived TIMP-1 for therapeutic targeting of neuronal and synaptic pathology in AD. In order to utilize a molecule as a therapeutic candidate it is essential to check how that molecule can be delivered to the brain. Although our route of injection (i.c.v.) suffices for a proof of-concept study to fully acknowledge TIMP-1’s wide-ranging beneficial effects in the brain in AD, in order to uphold its potential as a therapeutic candidate we discuss previous attempts that have been made *albeit* in different neurological disorders to deliver TIMP-1 in the brain. As discussed above, although an exogenous TIMP-1 injection may induce different autoregulatory loops to sustain its beneficial effects for a long duration, the half-life of injected TIMP-1 itself may be short (<4h) as previously reported^99^. Unfortunately, TIMP-1 cannot cross the BBB, and has a rapid clearance and low bioavailability. In order to overcome these hindrances, several attempts have been made and those successfully enhance TIMP-1’s bioavailability as well as its delivery across BBB following systemic injections. For instance, recombinant human TIMP-1 has been PEG(polyethylene glycol)ylated which increased its plasma half-life in mice from 1.1h to 28h^100^. Furthermore, loading TIMP-1 in poly(lactic-co-glycolic acid) nanoparticles coated with polysorbate 80^101^ and in magnetic nanoparticles^102^ increased its delivery across BBB. Interestingly, a single study shows that radiolabelled recombinant TIMP-1 injected intravenously in a rat model of cerebral ischemia had a long blood half-life of 42.2 h, was rapidly distributed in the brain tissue (within 0.5 h) and had slow elimination half-lives of 45.3 h and 39.2 h from the cortex and striatum respectively^103^. TIMP-1 being an endogenous molecule, to begin with, which means it had a minimum chance of showing toxicity as opposed to exogenous candidates, strongly emerge as a major beneficial player in AD through our study (both by its depletion as a cause and its restoration as a potential treatment). Together with consistent efforts being made by others to enhance its bioavailability and increase its delivery to the brain, TIMP-1 plausibly holds a strong clinical value for the treatment of AD in future.

## MATERIALS & METHODS

### Experimental Design

Multiple independent experiments were performed consisting of several biological replicates. Sample sizes and statistical analysis to calculate P values are given in detail in figure legends, in Table S1 and in the ‘Statistical analysis’ section. No statistical method was followed to pre determine the sample sizes, but sample sizes were kept similar to that in relevant publications in literature. Data distribution was not formally tested for normality, but rather assumed to be normal. Data were acquired and analyzed by multiple researchers, blind to the experimental conditions whenever experimentally feasible.

### Materials

Dulbecco’s modified Eagle’s medium (DMEM)-F12, Neurobasal medium, B27 supplement, Fetal Bovine Serum (FBS), goat serum, Penstrep antibiotic, Lipofectamine 2000, siCD63, scrambled siRNA (negative control), Sypro Ruby, Hoechst 33342, Prolong Gold Antifade with DAPI and Alexa Fluor were purchased from Thermo Fisher Scientific (MA, USA). Insulin, progesterone, putrescine, selenium, Apo-transferrin, poly-D-lysine (PDL), Di-methyl sulfoxide (DMSO), paraformaldehyde (PFA), bovine serum albumin (BSA), Triton-X 100, radioimmunoprecipitation assay (RIPA) buffer and 1,1,1,3,3,3 hexafluoro-2-propanol (HFIP) were purchased from Sigma Aldrich (St. Louis, USA). Cell culture dishes, plates, filter units and flasks were purchased from BD Falcon (Schaffhausen, Switzerland) and Corning (NY, USA). Aβ (1–42) was purchased from Alexotech (Vasterbotten County, Sweden). PVDF membrane and Percoll were purchased from GE Healthcare (Buckinghamshire, UK). Enhanced chemiluminescence (ECL) substrate and stripping buffer were purchased from Takara Bio (Kusatsu, Japan). Sodium dodecyl sulphate (SDS) was purchased from Merck (Darmstadt, Germany). Trypsin and Bovine Serum Albumin (BSA) were purchased from Sisco Research Laboratories Pvt. Ltd (Mumbai, India). All other fine chemicals were procured from standard local suppliers unless stated otherwise. List of antibodies used for this work, their dilutions for each type of application and sources are given in the table below.

**Table:** List of antibodies. WB=western blot, ICC=immunocytochemistry, IHC=immunohistochemistry, CST=Cell Signaling Technology, SCBT=SantaCruz Biotechnology

| **Antibody** | **Species raised** | **Application** | **Dilution** | **Source** | **Cat No.** |
| --- | --- | --- | --- | --- | --- |
| **phospho-Akt (Ser473)** | rabbit | WB | 1:500 | CST | 4058 |
| **phospho-Akt (Thr308)** | rabbit | WB | 1:500 | CST | 4056 |
| **Akt** | rabbit | WB | 1:1000 | CST | 9272 |
| **phospho-FOXO3a (Ser253)** | rabbit | WB | 1:500 | CST | 9466 |
| **FOXO3a** | rabbit | ICC | 1:100 | CST | 2497 |
|  |  | WB | 1:1000 |  |  |
| **BIM** | rabbit | WB | 1:1000 | Abcam | ab7888 |
| **PUMA** | rabbit | WB | 1:1000 | Novus | NBP1-76639 |
| **Bcl-2** | rabbit | WB | 1:1000 | CST | 2870 |
| **Bcl-xl** | rabbit | WB | 1:1000 | CST | 2762 |
| **Bax** | rabbit | WB | 1:1000 | CST | 2772 |
| **pH2Ax (Ser139)** | rabbit | IHC | 1:100 | CST | 9718 |
|  |  | WB | 1:1000 |  |  |
| **Cleaved Caspase 3 (Asp175)** | rabbit | WB | 1:1000 | CST | 9661 |
| **ATG5** | rabbit | WB | 1:1000 | Novus | NB110-53818 |
| **p62** | rabbit | WB | 1:1000 | Novus | NBP1-42821 |
|  |  | ICC | 1:100 |  |  |
|  |  | IHC | 1:100 |  |  |
| **LC3B** | rabbit | WB | 1:1000 | Novus | NB100-2220 |
|  |  | ICC | 1:100 |  |  |
|  |  | IHC | 1:100 |  |  |
| **LAMP-1** | rabbit | WB | 1:1000 | Abcam | ab24170 |
|  |  | ICC | 1:100 |  |  |
|  |  | IHC | 1:100 |  |  |
| **TIMP-1** | mouse | WB | 1:750 | SCBT | sc-21734 |
|  | rabbit | ICC | 1:100 | Millipore | AB770 |
|  |  | IHC | 1:100 |  |  |
| **CD63** | mouse | WB | 1:750 | SCBT | sc-5275 |
|  |  | ICC | 1:100 |  |  |
| **Aβ** | rabbit | IHC | 1:100 | Abcam | ab201060 |
| **pULK1(Ser757)** | rabbit | IHC | 1:100 | CST | 6888 |
| **PSD-95** | rabbit | IHC | 1:100 | ThermoFisher Scientific | 51-6900 |
|  |  | WB | 1:1000 |  |  |
| **SNAP-25** | mouse | IHC | 1:100 | SCBT | sc-20038 |
|  |  | WB | 1:1000 |  |  |
| **Synaptophysin1** | mouse | WB | 1:2000 | Synaptic Systems | 101011 |
| **Homer1** | mouse | WB | 1:2000 | Synaptic Systems | 160011 |
| **Actin** | rabbit | WB | 1:10000 | Sigma Aldrich | A3854 |
| **GAPDH** | rabbit | WB | 1:1000 | CST | 2118 |
| **phospho-Cofilin(Ser3)** | rabbit | WB | 1:500 | CST | 3311 |
| **Cofilin** | rabbit | WB | 1:1000 | CST | 3318 |
| **phospho-GSK3β (Ser9)** | rabbit | WB | 1:1000 | CST | 5558 |
| **GSK3β** | rabbit | WB | 1:2000 | CST | 9315 |
| **Pro-BDNF** | mouse | WB | 1:250 | SCBT | sc-65514 |
| **TrkB** | mouse | WB | 1:500 | ABclonal | A21227 |
| **pERK1/2 (Thr202/Tyr204)** | rabbit | WB | 1:1000 | CST | 4377 |
| **ERK2** | rabbit | WB | 1:1000 | SCBT | sc-153 |
| **pCREB (Ser133)** | rabbit | WB | 1:500 | CST | 9198 |
| **CREB** | rabbit | WB | 1:500 | CST | 4820 |
| **Anti-rabbit IgG-**<br>**HRP inked** | Goat | WB | 1:4000 | CST | 7074 |
| **Anti-mouse IgG-**<br>**HRP inked** | Horse | WB | 1:4000 | CST | 7076 |

### Primary Cortical Neuron Cultures

Primary cortical neuron culture was performed following the method described earlier^32^. In short, the neocortex regions of E16-18 (sex not determined) of Sprague Dawley rats were dissected out under a dissection microscope in serum free medium (DMEM/F12 [1:1]) supplemented with 6 mg/ml D-glucose, 100 μg/ml transferrin, 25 μg/ml insulin, 20 nM progesterone, 60 μM putrescine, 30 nM selenium) along with penstrep. Following gentle trituration, the single cell suspension was seeded at 1.2 million in 2 ml per well of a previously PDL-coated 6-well tissue culture plate. After one day *in vitro* (1 DIV), medium was changed with fresh Neurobasal medium supplemented with B27. On 4 DIV, half of the used medium was replaced with fresh neurobasal plus B27 medium. Following six days of plating the neurons were treated as per experimental conditions in fresh neurobasal plus B27 medium.

### Preparation of oligomeric Aβ

HFIP solvent (100%) was added to lyophilized Aβ_1-42_ peptide to a final concentration of 1 mM and centrifuged in a speed vac (Eppendorf, Hamburg, Germany). The peptide pellet was re suspended in DMSO at a 5 mM concentration and sonicated in a 37⁰C water bath for 10 min. The peptide solution was aliquoted and stored at -80°C for later use. Before treatment, each aliquot was diluted in phosphate buffer saline (PBS: NaCl 137 mM, KCL 2.7 mM, Na_2_HPO_4_ 10 mM, KH_2_PO_4_ 2 mM, pH 7.4) and SDS (0.2%) to a concentration of 400 μM and incubated for 18-24 h at 37⁰C. Next, the suspension was further diluted with PBS to a final concentration of 100 μM and again incubated for 18-24 h at 37⁰C. The Aβ suspension was analyzed by SDS-polyacrylamide gel electrophoresis (SDS-PAGE) followed by Sypro Ruby protein gel staining to check for oligomeric species.

### Treatments of cells

Cortical neurons were treated with either 1.5 µM Aβ or rat recombinant TIMP-1 peptide (R&D systems, Cat# 580-RT-010, Minneapolis, USA) at 100 ng/ml^32^ or a combination of both for 16 h as shown in figures.

### Transfection studies

Primary neuron cultures were treated with Silencer Select siCD63 (s130868, Thermofisher Scientific, Waltham, MA, USA) or Silencer Select Negative control No. 1 siRNA (4390843, Thermofisher Scientific, Waltham, MA USA) on Day-4 following plating according to the manufacturer’s protocol at 100 pmole/well (50 nM) of a 6-well tissue culture plate with the help of Lipofectamin 2000. Successful knockdown was verified following 72 h of incubation by western blot for CD63 in siCD63 transfected cells versus Silencer Select Negative control No. 1 siRNA and untreated control cells. On Day-7 following plating, the cells were treated according to the indicated experimental paradigm.

### Immunocytochemistry

Neurons on PDL-coated glass coverslips were fixed in 4% PFA in PBS for 15-20 min at room temperature. Following a PBS wash, the fixed cells were permeabilized and blocked with 3% goat serum in 0.3% Triton X-100 in PBS for 1-2 h. Next, cells were incubated with primary antibody overnight at 4°C. Following the primary antibody incubation, cells were washed and incubated with specific species-matched secondary antibody tagged with Alexa fluor 546/488 for 1-2 h at room temperature. Nuclei staining were performed using Hoechst 33342 in PBS at 2 µg/ml for 30 min in the dark. Cells were then mounted on glass slides using Prolong Gold antifade with DAPI. Images were taken in Leica Sp8 STED confocal microscope (Wetzlar, Germany) at specific magnifications as mentioned in individual experiments. The fluorescence intensities for FOXO3a in the nucleus and whole cell were calculated separately by NIH-ImageJ software. Fluorescence intensity for FOXO3a in the cytoplasm was calculated by subtracting the nuclear FOXO3a from total cell FOXO3a. The ratio of nuclear FOXO3a to cytosolic FOXO3a under each condition is presented to indicate the translocation. In the case of colocalization studies, Pearson’s correlation coefficient was calculated from images taken at 60x magnification using Leica Sp8 software. Integrated density of staining, area of individual cells and the background fluorescence of individual images in different experimental conditions was derived using NIH-ImageJ software. The corrected total cell fluorescence (CTCF) was calculated as follows, CTCF = Integrated density – (area of selected cell x mean fluorescence of background readings).

### Preparation of cell or tissue lysate

Following treatment, cells were washed and scraped in PBS and harvested by centrifugation at 800 g at 4⁰C for 5 min. In case of tissues, cortex and hippocampus from mouse brain were micro dissected, the tissue was immediately collected in RIPA buffer along with phosphatase inhibitor (Roche, Basel, Switzerland) and protease inhibitors (Takara Bio, Kusatsu, Japan) mixes. Alternatively in case of culture, cells were lysed with RIPA buffer (Sigma-Aldrich, St. Louis, USA) containing with phosphatase inhibitor and protease inhibitor. For both tissue and cell pellets, each tube contents was sonicated by 2-3 pulses on ice to breakdown genetic material and incubated for 30 min on ice with intermittent vortexing. The lysed cells/tissue was then mixed with 5x sample buffer so that the final contents are at 1x sample buffer concentration. The protein concentration was estimated by the Lowry method.

### Western blotting

Fifty µg of protein from each treatment condition was run in SDS-PAGE. The resolved proteins were then transferred to PVDF membrane. The membranes were next blocked with 5% BSA for 1 h at room temperature, following which the membranes were incubated with primary antibodies overnight at 4⁰C. Membranes were washed with TBST [1.5 M NaCl, 1 M Tris (pH 7.5), 0.1% Tween20] followed by incubation with HRP-conjugated suitable secondary antibody for 1 h at room temperature. Three more TBST washes were given to remove unbound antibodies. Protein bands were detected on ChemiDoc MP imaging system (Bio-Rad laboratories, California, USA) using ECL reagents. Densitometric quantification of each band was done using NIH ImageJ software (NIH, Bethesda, MD, USA), normalized with loading control (either Actin or GAPDH as mentioned) and fold-changes were compared with the untreated control.

### Animal housing and care

Male C57BL6 (wild-type or WT) or 5xFAD mice were housed at a maximum of six per cage in standard individually ventilated cages (IVC) with proper bedding material. Mice were maintained by trained caretakers, checked daily and kept in a temperature-controlled room (24 ± 2°C), with a 12 to 12 h light-dark cycle, humidity (60 ± 5%) and access to food and water ad libitum in the animal house of the CSIR-Indian Institute of Chemical Biology, Kolkata. For age-wise comparisons, 5xFAD mice of different ages as defined later were procured along with age-matched WT mice. For all other animal experiments, 6-month-old 5xFAD and age-matched WT mice from different litter groups were used for treatment. All the studies were conducted in accordance with the guidelines formulated by the Committee for the Purpose of Control and Supervision of Experiments on Animals (Department of Animal Husbandry and Dairying, Ministry of Fisheries, Animal Husbandry and Dairying, Govt. Of India), with approval from the Institutional Animal Ethics Committee (IAEC) on the required care and use of animals in laboratory research.

### Stereotaxic procedures

For mouse, a combination of ketamine (100 mg/kg i.p.) and xylazine (10 mg/kg i.p.) in 0.9% normal saline was used for anesthetization (i.p.) for a short duration for stereotaxic surgery. Individual mice were placed in a suitable stereotaxic frame (Stoelting, MO, USA) with the incision bar initially positioned at the Bregma point. The body temperature of the animal was maintained at 37°C using a homeothermic blanket (Harvard apparatus, MA, USA). Recombinant TIMP-1 (5 ng) diluted in artificial cerebrospinal fluid (aCSF) in a total volume of 5 µl was injected intracerebroventricularly (i.c.v.) into the ventricle (lateral, 0 cm from the bregma; anteroposterior, -1.6 mm from bregma; dorsoventral, -2.2 mm from bregma). The dose of TIMP-1 was standardized in our previous study^11^ and can be correlated to the concentration of TIMP-1 detected in the CSF of healthy individuals (human adults) ranging from 60 ng/ml to 160 ng/ml^16^ calculated with respect to the average CSF volume in mice (around 40 µl) where the intracerebroventricularly injected TIMP-1 gets distributed. The flow rate was kept at 0.5 µl/min using a worker bee syringe pump (BAS, West Lafayette, USA). The peptide solution was allowed to diffuse for an additional 10 min following actual injection duration to prevent backflow.

5xFAD mice were randomly divided into three groups to receive treatments – 5xFAD, 5xFAD+aCSF, 5xFAD+TIMP-1 (5 ng). WT mice were randomly divided into two groups – one without any treatment and another with TIMP-1 injection. WT+TIMP-1 group was not included in the behavioral tests (since we did not find any change with TIMP-1 only icv injection in rat behaviors in our earlier study^11^) but was included for synaptosome and electrophysiological studies.

### Behavioral assessments

Behavioral tests were started on Day-8 following TIMP-1 infusion and performed in a blind-folded manner.

#### Open-field test (OFT)

Open-field test (OFT) apparatus (IR Actimeter, Panlab, Barcelona, Spain) consists of an open field Plexiglas arena with photocell emitters and receptors positioned at equal distance along the perimeter of the arena (278×236×300 mm). These emitters and receptors create an x-y grid of invisible infrared rays. Mouse moving across the arena makes beam breaks that were recorded for 10 min and the animal trajectories were analyzed by ActiTrack software (Panlab, Barcelona, Spain). The arena was divided into two zones; a square area occupying one-third of the total arena area in the center was considered as the ‘Inner zone’ and the rest the ‘Outer zone’. The total distance travelled (cm), the speed of the mouse (cm/sec) and the time spent in the outer zone (periphery) and that in the inner zone (center) in seconds were recorded, where the latter is generally utilized to measure anxiety.

#### Elevated plus maze (EPM) test

The EPM test for long-term memory was carried out according to the standard protocol as described earlier^32^. Briefly, in the two-day test, on day one the animal was placed at the edge of one of the open arms of plus maze with its face away from the center for exploration for 5 min. On day two, the mouse was placed similarly on to the open arm for 5 min. The transfer latency to the closed arm on day one was recorded as initial transfer latency (ITL) and on day two was recorded as retention transfer latency (RTL).

#### Novel Object Recognition (NOR) test

In this two-day test for hippocampus-dependent recognition memory, a mouse was allowed to explore a rectangular box (50 cm x 30 cm) with two identical (color and dimensions) objects placed at parallel corners for 5 min on Day-1. On Day-2, one of the objects was replaced with an object of different color and shape and the mouse was returned to the box for 5 min. The time the mouse spent with each of the two objects on Day-2 were noted and presented as exploration time (sec). The Discrimination index (DI) and preference index (PI) were derived according to the formulas – DI = (TN-TF)/(TN+TF), PI = TN/(TN+TF)*100 from the Day-2 data where TF is time spent with the familiar object and TN is time spent with the novel object.

#### Fear Conditioning Test

Fear conditioning tests are routinely utilized to study long-term associative learning processes involving amygdala, hippocampus and specific parts of the cortex. Each mouse was placed in a sound-attenuating chamber with a stainless-steel grid floor that delivers a foot-shock (Panlab, Barcelona, Spain) and is connected to a recorder that records the freezing behavior. Rodents express their fear towards an aversive stimulus with a complete loss of movement except for breathing termed ‘freezing’. Animals were handled once per day for three days and habituated in the experimentation room before the start of the actual experiment. Only one animal was allowed in the experimentation room at a time. Both context-dependent and cue-dependent fear conditioning tests were performed as described earlier^32^ with modifications as mentioned below.

#### Contextual fear conditioning test

On the day of training, the animal was placed in the chamber and allowed to acclimatize for a 2 min period. After acclimatization, the mouse was exposed to four sessions of an unconditional stimulus (each session 89 sec pre-shock interval; 0.4 mA, 1 sec foot-shock) co-terminating with a 2 min period without stimuli (10 min total). Context dependent fear memory retention was tested on day two when the mouse was returned to the same environment but this time without foot shock. Freezing behavior was calculated by PACKWIN 2.0 software (Panlab, Barcelona, Spain) during an 8 min run time.

#### Cue-dependent fear conditioning test

On the day of training, each mouse was initially exposed to the chamber for a 2 min period for acclimatization. Next, four 29-sec sessions of light and sound (80 dB, 2500 Hz) cues co terminating with a foot shock (0.4 mA, 1 sec for mouse) were delivered with an inter-trial interval of 1 min, following which the mouse was given 2 min of rest (10 min in total) before it was removed from the chamber. On day two, the context of the chamber was altered by attaching white sheets to the chamber walls and the floor and the mouse was returned to this new environment with the same sessions of cues but without foot-shock. Cue-dependent fear memory retention was assessed by recording freezing behavior during the 8 min test period by PACKWIN 2.0 software (Panlab, Barcelona, Spain).

### Immunohistochemistry

After 14 days of TIMP-1 infusion, each mouse was transcardially perfused under anesthesia with urethane [1g/kg body weight (MP Biomedicals, lllkirch, France) in 0.9 % NaCl solution (i.p.)] with chilled 100 mM PBS (pH 7.4). Perfusion was continued with 4% (w/v) PFA solution in PBS, following which the mouse was decapitated and the brain was isolated in chilled 4% PFA solution for overnight fixation at 4°C and then transferred to a 30% (w/v) sucrose solution in PBS for cryopreservation at 4°C. Coronal sections (20 µm thick) were obtained using a cryotome (Thermo Shandon, Pittsburg, USA) and collected on gelatine-coated slides for immunohistological staining. Selected brain sections were washed with PBS thrice followed by permeabilization with PBS + 0.4% Triton-X for 40 min. Next, the sections were washed with PBS + 0.1% Triton-X (PBST) three times, 5 min each. The sections were then blocked with 4% BSA in PBST for 1 h. Following blocking, the sections were incubated with primary antibodies diluted in the blocking solution for 48 h at 4°C. Next, PBST washes were given to the tissue slices to remove unbound antibodies and incubated with appropriate Alexa fluor-tagged secondary antibodies diluted in the blocking solution at RT for 2 h. Afterwards, sections were again washed in PBST and incubated with Hoechst 33342 in PBS at 2 µg/ml for 30 min at 37°C. Further PBS washes were given to the sections and mounted with Prolong Gold Antifade with DAPI reagent on glass slides for microscopic analysis in Leica Sp8 STED confocal microscope (Wetzlar, Germany).

Aβ1-42 plaques were evaluated by calculating the percentage of plaque area and plaque number by ImageJ software from 40x images of the hippocampus taken in Leica Sp8 STED confocal microscope (Wetzlar, Germany). For SNAP25, PSD95, pULK1, p62, LC3B and LAMP1, CTCF = Integrated density – (area of selected cell x mean fluorescence of background readings) was calculated from cells in the cortices and hippocampi of mouse brain sections with ImageJ software from 40x images taken in Leica Sp8 STED confocal microscope (Wetzlar, Germany). TIMP-1 count denotes the number of TIMP-1 antibody-stained puncta in the hippocampal-cortical regions from 20x images of mouse brain sections taken under a confocal microscope. The number of pH2Ax positive nuclei and the total number of nuclei in the specific area was counted by ImageJ software from 100x images taken in the hippocampal and cortical regions by a confocal microscope and the percentage was calculated.

### Terminal deoxynucleotidyl transferase dUTP nick end labeling (TUNEL) assay

TUNEL assays were performed to detect apoptotic cell death using Clonetech ApoAlert DNA Fragmentation kit (Takara, Kusatsu, Shiga, Japan) according to the manufacturer’s protocol. Fixed tissue sections were hydrated with PBS and incubated with Proteinase K solution (20 µg/ml, diluted from the stock provided in the kit) for 20 min. The reaction was terminated by washing with PBS, followed by addition of 4% PFA in PBS to the sections. Following PBS washes, tissue sections were equilibrated in equilibration buffer at RT for 10 min. The slices were then incubated in a solution of nucleotide mix and Tdt enzyme for 1 h at 37°C in the dark. The tailing reaction was stopped by addition of 2XSSC buffer (provided in the kit) at RT for 15 min. Next, the sections were rinsed again in PBS and incubated with Hoechst solution for 30 min at 37°C, washed with PBS and mounted with Prolong Gold Antifade with DAPI for microscopic analysis. The percentage of apoptotic cells were calculated by dividing the number of TUNEL-positive cells by the total number of cells (Hoechst-stained nuclei) multiplied by 100.

### Isolation of synaptosomes

Synaptosomes were isolated from adult mouse brain by the method described earlier with some modifications^40^, 14 days following TIMP-1 infusion. Briefly, the freshly dissected mouse brain tissue (hippocampus or cortex), pooled from six to seven mice per experimental group, was homogenized in an isotonic sucrose gradient buffer (diluted to 1x from a stock of 4x Gradient buffer 1.28 M sucrose, 4 mM EDTA, 20 mM Tris, pH 7.4). The homogenate was then centrifuged at 1000g for 10 min at 4°C to separate supernatant and debris. The supernatant was loaded onto a discontinuous Percoll gradient composed of three layers – 3% (vol/vol), 10% (vol/vol) and 23% (vol/vol) percoll. The gradient was then centrifuged at 31,000 g for 5 min at 4°C (excluding acceleration and deceleration time). The synaptosomes separate as a layer between the 10% and 23% percoll layers. The synaptosome layer was collected, diluted in aCSF and centrifuged to remove percoll contamination. The final synaptosomal pellets were re-suspended in 1x sample buffer before protein estimation for SDS-PAGE run. Alternatively, for determination of Fibrillar/Globular (F/G) actin content ratio, the synaptosomal pellet was further fractionated into the cytoskeleton and cytoplasmic fractions. Accordingly, the synaptosomal pellet was resuspended in a Mg^2+-^containing buffer (50 mM Tris, 2 mM MgCl2, pH 7.4) and vortexed while maintaining the suspension on ice. Next, TritonX-100 was added to the suspension at a final concentration of 0.5% and kept on ice for 20-30 min. Then, it was centrifuged at 12,000 g for 10 min at 4°C. The pellet thus obtained is the cytoskeleton (CSK) fraction containing F actin while the supernatant (SUP) contains all the cytosolic proteins including G actin. Each of the fractions was resuspended in 1x sample buffer for western blot analysis.

### Golgi-cox impregnation and staining procedure

Golgi-cox staining was performed according to the protocol described previously^104^. Fourteen days following TIMP-1 infusion, the brain of the mouse was taken out and immediately submerged in the impregnation solution a mixture of five parts by volume of Solution A (5% potassium dichromate in distilled water (DW)), five parts by volume of Solution B (5% mercuric chloride in DW), four parts by volume of Solution C (5% potassium chromate in DW) and ten parts by volume of DW, and kept in dark at RT. The solution was changed after 24 h with fresh impregnation solution and incubated for two weeks. Next, the brain was transferred to Solution D, a cryoprotectant solution prepared by mixing 20 g of sucrose and 15 ml glycerol in 100 ml of DW and kept at 4°C in the dark for 24 h following which fresh cryoprotectant solution was added. A period of 2-3 days was given for the brain to sink to the bottom. Sectioning was done at -25°C in a cryotome (Thermo Fisher Scientific, MA, USA) and 100 µm thick slices were collected on gelatin-coated glass slides. The sections were allowed to dry for some time, washed twice with DW and stained with 20% ammonium solution for 10 min in dark. Following two more washes with DW, sections were dehydrated through 50%, 70%, 95% and 100% (twice) ethanol for 5 min in each. The sections were removed of fat by xyline wash, twice for 5 min each. The sections were mounted in DPX reagent, covered with a coverslip and allowed to dry at room temperature for 3-4 days in the dark before bright-field imaging in Leica Sp8 STED confocal microscope (Wetzlar, Germany). For measuring spine density, distal dendrites with similar diameters were imaged in the cortex or stratum radiatum of the CA1 hippocampus, magnified in Adobe Photoshop (San Jose, California, USA) and spines counted on printed copies, as described earlier^105^. Any small protrusion on a dendrite was considered a spine. Spine density was calculated from many distal dendrites amounting for a total length of 1200-2000 µm (40-80 dendritic stretches) per experimental group.

### Electrophysiology

Fourteen days following TIMP-1 injection, mice were anesthetized in a 5% gaseous isoflurane chamber (Aerrane, Baxter Healthcare Corporation; Fluovac, Harvard Apparatus, USA) and perfused transcardially at a speed of 45 ml/min using a peristaltic pump (Cole Parmer, USA) with cold (7-8°C) aCSF [NaCl (118 mM), KCl (2.5 mM), NaH_2_PO_4_ (1.5 mM), NaHCO_3_ (25 mM), C_6_H_12_O_6_ (25 mM), MgCl_2_ (1.3 mM) and CaCl_2_ (2.2 mM), adjusted to a pH of 7.4 with an osmolarity of 310-320 mOsm/L] which was continuously bubbled with carbogen (5% CO_2_ and 95% O_2_, Chemix Special Gases and Equipments, India). The brain was immediately removed from the skull and submerged in ice-cold carbogenated aCSF. The hemispheres were separated and 400μm thick para-sagittal slices were taken using a vibratome (Leica VT, 1200S, Leica Biosystems, Germany) with a blade speed of 0.14 mm/s and amplitude 1.20 mm. The slices were transferred and kept in a brain slice keeper (BSK4-Quad, Automate Scientific Inc CA, USA) in carbogenated aCSF at RT. For extracellular Field Excitatory Postsynaptic Potential (fEPSP) recordings, after 2 h incubation at RT, slices were transferred to a recording chamber (Warner instruments, USA), perfused continuously with carbogen-bubbled aCSF at RT with the help of a Minipuls Evolution peristaltic pump (Gilson Inc USA). The flow rate was maintained at 2 ml/min. Under observation in a stereo zoom microscope (Olympus, Japan), platinum-iridium cluster bipolar electrodes (Frederick Haer Company Inc, ME, USA) were placed to stimulate Schaffer collaterals that synapse with CA1 pyramidal neurons. Borosilicate glass microelectrode (Sutter instruments, USA) filled with 2M NaCl was used as the recording electrode and was placed in the stratum radiatum layer of the CA1 region of dorsal hippocampus, and recordings were obtained under immersed conditions. fEPSPs were induced by injecting 0.1-ms width square wave pulses using a stimulus isolator in a constant current mode that produced fEPSP with a typical stimulus artefact, fibre volley and a negative deflection. fEPSPs were induced at a test stimulus frequency of 0.033Hz during the whole experimental duration. A stable baseline of minimum 20 min was recorded before proceeding with the evaluation of other parameters. The studies conducted sequentially were as follows:

i. **Input-output (I-O) relationship** was measured by increasing the input current strength (by 20 µA increments) until population spike was observed. The stimulus current strength for the test was kept at a value that evoked a 40% of the maximum fEPSP slope value.
ii. **Paired-pulse facilitation (PPF)** was determined by supplying two pulses in close succession with inter-stimulus intervals (ISI) ranging from 30-300 ms with an increment of 30 ms.The paired-pulse ratio (PPR) was calculated by dividing the slope of fEPSP2 by fEPSP1.
iii. Following 20 min of baseline recording, **post-tetanic potentiation** (PTP), **short term potentiation** (STP) and **long-term potentiation** (LTP) were evoked using single theta burst stimulation (TBS, 4 pulses/burst at 100 Hz, 20 bursts with 200-ms interval). Recordings were carried out up to 60 min following TBS induction. Only slices that displayed stable responses for a minimum of 60 min post TBS were considered for analysis of PTP, STP and LTP. The average slopes of fEPSPs normalized to baseline measured for the first 3 min (PTP), from 21^st^-30^th^ min (for STP) and 41^st^-60^th^ min (for LTP) post TBS were used for statistical comparisons.

The fEPSP responses were amplified using a differential amplifier (A-M systems, USA) and digitized with the help of a digitizer (National Instruments, USA). A noise eliminator (Humbug, Quest scientific) was utilized to eliminate the effect of electrical interference on the recordings. The data were acquired and analyzed using WinLTP® v2.3 software (WinLTP Ltd, UK).

### Statistical Analysis

All data were analyzed and graphs were created by using Prism software (v8.4.2 (679), GraphPad Inc, USA). For analysis of significant difference between only two groups two-tailed unpaired student’s t-test was used. For comparing more than two experimental groups, one-way ANOVA analysis followed by Tukey’s multiple comparison post hoc test was used in all the experiments except as stated otherwise. However, in the case of the input-output curve and PPF curve, two way ANOVA followed by Tukey’s multiple comparison post hoc analysis was used to calculate the effect of TIMP-1. All data are presented as mean ± standard error of mean (s.e.m.), and P<0.05 was considered to be statistically significant.

## DATA AVAILIBILITY

This study includes no data deposited in external repositories.

## ACKNOWLEDGEMENTS

We, the authors, would like to extend our gratitude to Prof. Lloyd Greene, Department of Cell Biology and Pathology, Columbia University, New York for reading this manuscript, correcting errors relating to grammar, critical comments, and helpful suggestions. We would like to thank Dr. P. K. Sarkar for critical reading of this manuscript and helpful discussions. We would like to thank Ms. Sammani Sarkar for her help in preparing the artwork of the manuscript. We would also like to thank Mr. Sounak Bhattacharya for his technical assistance in confocal microscopy. The work was supported by one of the 12^th^ Five Year Plan Projects, miND (BSC0115) of CSIR, Govt. of India.

## AUTHOR CONTRIBUTIONS

Conceptualization, S.S. and S.C.B.; Methodology, S.S., K.G. and R.K.P.; Validation, S.S. and K.G.; Formal analysis, S.S. and K.G.; Investigation, S.S., K.G. and R.K.P., Writing – Original Draft, S.S.; Writing – Review and Editing, S.S., S.C.B., K.G. and B.N.S.; Visualization, S.S. and S.C.B.; Funding acquisition, S.C.B.; Resources, S.C.B. and B.N.S.; Supervision, S.C.B. and B.N.S.

## SUPPLEMENTARY INFORMATION

### Supplementary Figures

**Figure S1.**
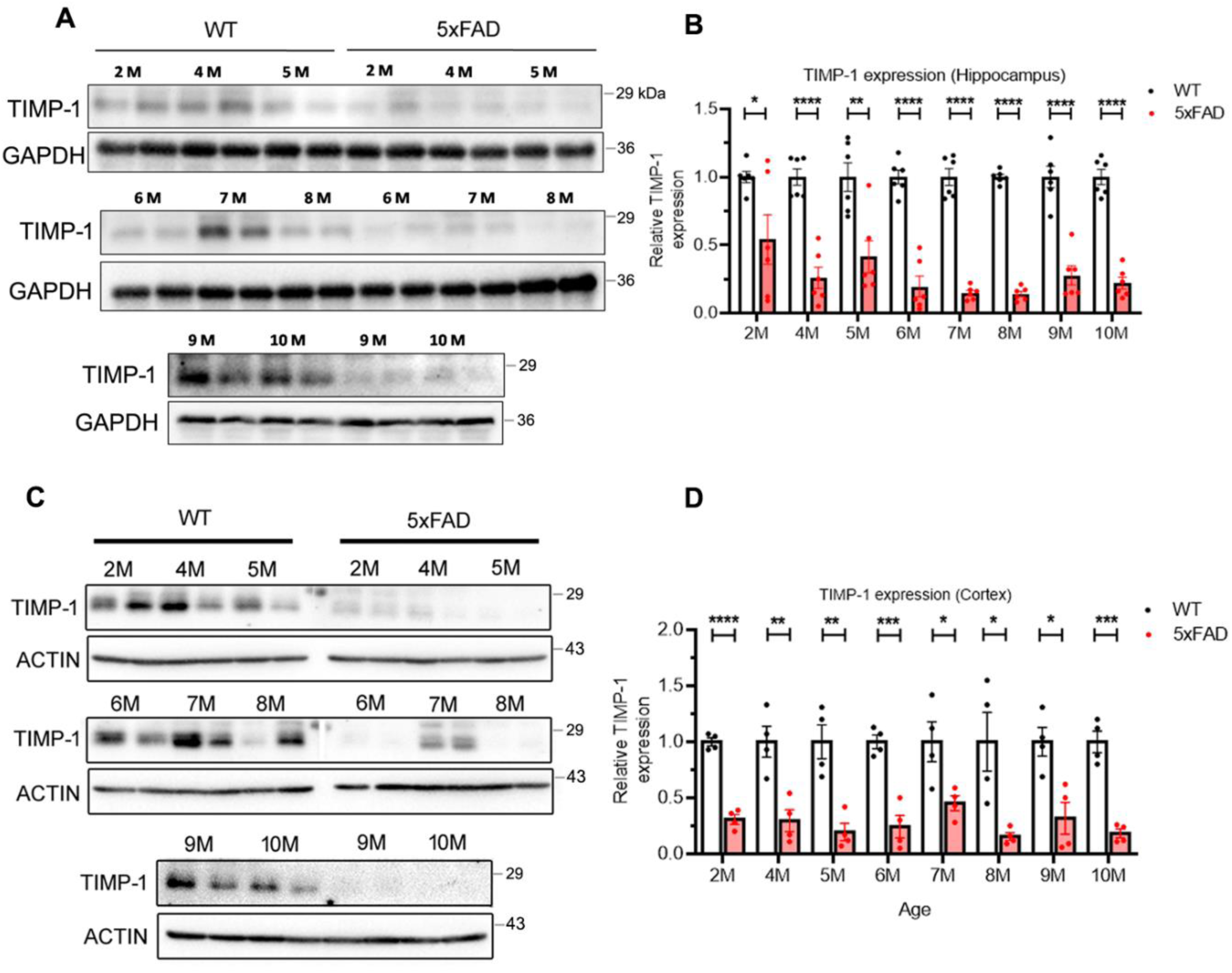
TIMP-1 expression in 5xFAD mouse is lower than WT mouse across age. Whole tissue lysates were obtained from WT (C57BL6) and 5xFAD male mouse hippocampi (**A-B**) and cortices (**C-D**) of different ages as indicated – 2 months (2M), 4M to 10M. **A,C**, Representative immunoblots for TIMP-1 from hippocampal (**A**) and cortical (**C,** ladder was run in the middle for 2M-5M and 6M-8M blots and cut through the ladder for convenience during blot development) whole tissues where two consecutive lanes indicate protein levels in two separate animals of each age group. **B,D,** Densitometric analysis for TIMP-1 normalized with GAPDH or Actin as the loading control from hippocampal (**B**) and cortical (**D**) blots, showing relative TIMP-1 expression in 5xFAD mice w.r.t. WT mice. For N=6 mice/group for **B** and N= 4 mice/group for **D**; two-tailed unpaired t-test, mean±s.e.m.*P<0.05, **P<0.01, ***P<0.001 and ****P<0.0001.

**Figure S2:**
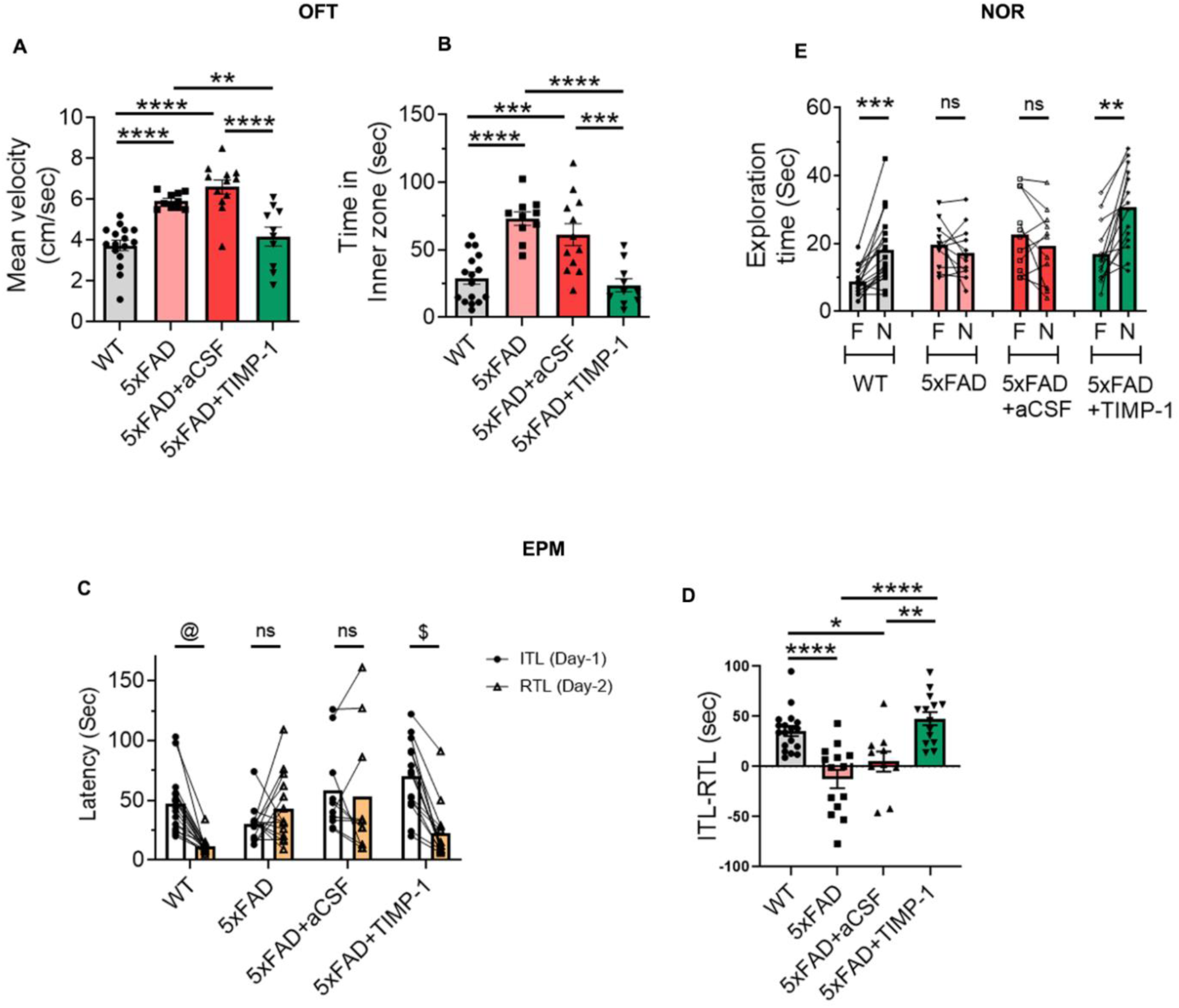
TIMP-1 injection improves cognitive performance of 5xFAD mice in behavioral tests. **A-B,** In open-field test (OFT), the mean velocity of the mouse in cm/sec (**A**) and the time spent in the inner zone in sec (**B**) of the OFT arena during 10 min duration. One-way ANOVA, Tukey’s post hoc, mean±s.e.m. **C-D**, In Elevated Plus-maze (EPM) test, latency (sec) to the closed arms on Day-1 (Initial transfer latency, ITL) and on Day-2 (Retention transfer latency, RTL), tracked for each mouse (**C**) and their difference ITL-RTL in sec for each group (**D**). Two-tailed unpaired t-test for **C** (@P<0.0001, $P<0.001) and One-way ANOVA, Tukey’s post hoc for **D**; mean±s.e.m. **E**, Day-2 results of Novel Object Recognition (NOR) test showing Exploration time (sec) with Familiar (F) versus Novel (N) object within each group for the 5 min duration of the test where the each connecting line tracks the exploration time for each mouse between F and N. Two-tailed unpaired t-test, mean±s.e.m. *P<0.05, **P<0.01, ***P<0.001 and ****P<0.0001; ns, not significant. Number of animals in each test and exact P values are given in Table S1.

**Figure S3.**
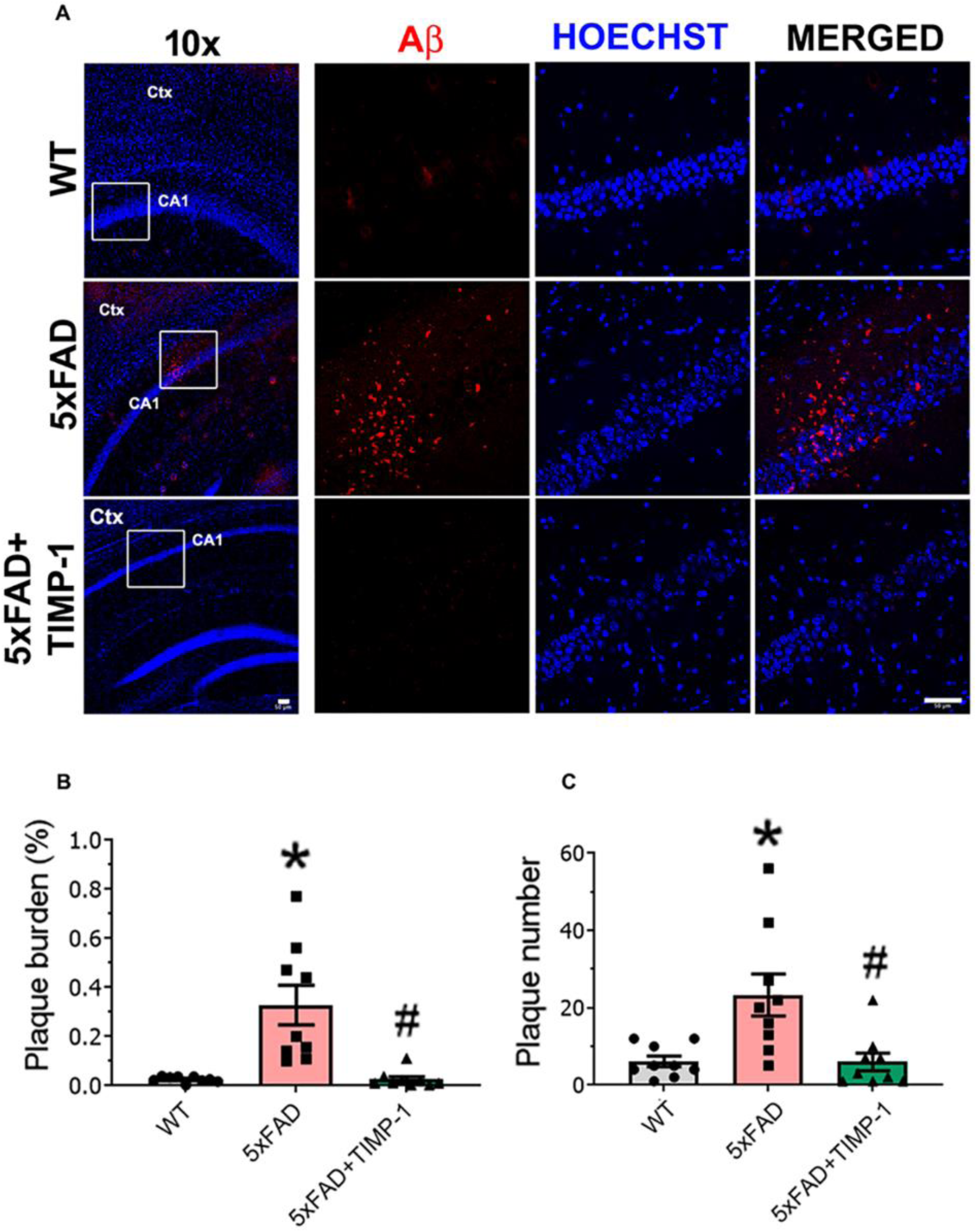
TIMP-1 injection in 5xFAD mice reduces Aβ plaques. **A**, Representative immunofluorescence images for Aβ_1-42_ deposits (red) in WT, 5xFAD and 5xFAD+TIMP-1 mouse brains in the hippocampal CA1 region, 14 days post TIMP-1 (5 ng) treatment. Nuclei stained by Hoechst (blue). Left panel shows 10x images (scale bar – 50 µm) where white box indicates region-of-interest in the CA1 region that were zoomed-in at 40x (scale bar – 50 µm). **B-C**, Plaque burden percentage (**B**, *P=0.0054 versus WT, ^#^P=0.0051 versus 5xFAD) and plaque number (**C**, *P=0.0004 versus WT, ^#^P=0.0004 versus 5xFAD) were calculated from area and number of Aβ puncta respectively. N=3 mice/group, n=3 slices/mouse brain. One-way ANOVA, Tukey’s post hoc, mean±s.e.m.

**Figure S4.**
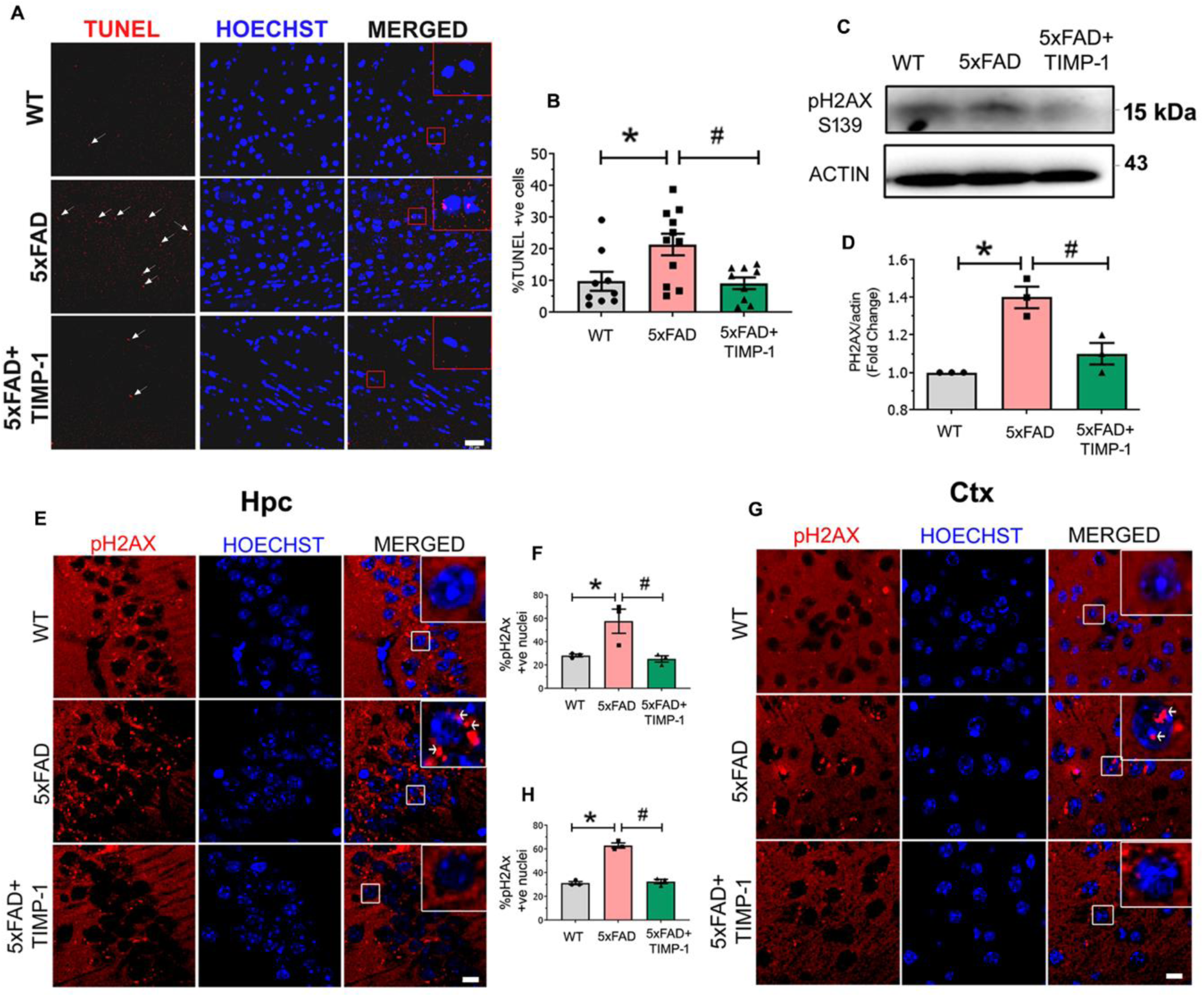
TIMP-1 restricts apoptosis in 5xFAD mice. **A**, Representative immunofluorescence images following TUNEL assay in 5xFAD mice sections showing apoptotic cells (red puncta highlighted with white arrow) colocalized with the nuclei (blue) imaged at 40x (scale bar – 25 µm), alongside WT and 5xFAD+TIMP-1 images. In merged image panel, the red box demarcates the nuclei zoomed-in for better viewing of the colocalization in the inset. **B,** Percentage of TUNEL positive cells for N = 3 mice/group, n=3-4 slices/mouse brain, 250 cells/slice at 20x. One-way ANOVA, Tukey’s post hoc, mean±s.e.m.; *p=0.0231, ^#^p=0.0162. **C**, Representative western blots for pH2Ax(S139) from hippocampal tissue along with respective actin. **D,** Densitometric analyses for pH2Ax showing fold change w.r.t. WT. N=3 mice/group; One-way ANOVA, Tukey’s post hoc, mean±s.e.m.; *P<0.05, ^#^P<0.05. **E,G**, Representative immunofluorescence images for pH2ax (red) and nuclei (blue) in the hippocampus (**E**) and in cortex (**G**); scale bar 10 µm. White box shown in the merged panel demarcates a single nucleus zoomed for higher magnification viewing (inset). **F,H,** Percentage of pH2Ax positive nuclei in hippocampus (**F**, *p=0.0367, #p=0.0246) and cortex (**H**, *p<0.0001, #p <0.0001), N=3 mice/group. One-way ANOVA, Tukey’s post hoc, mean±s.e.m.

**Figure S5:**
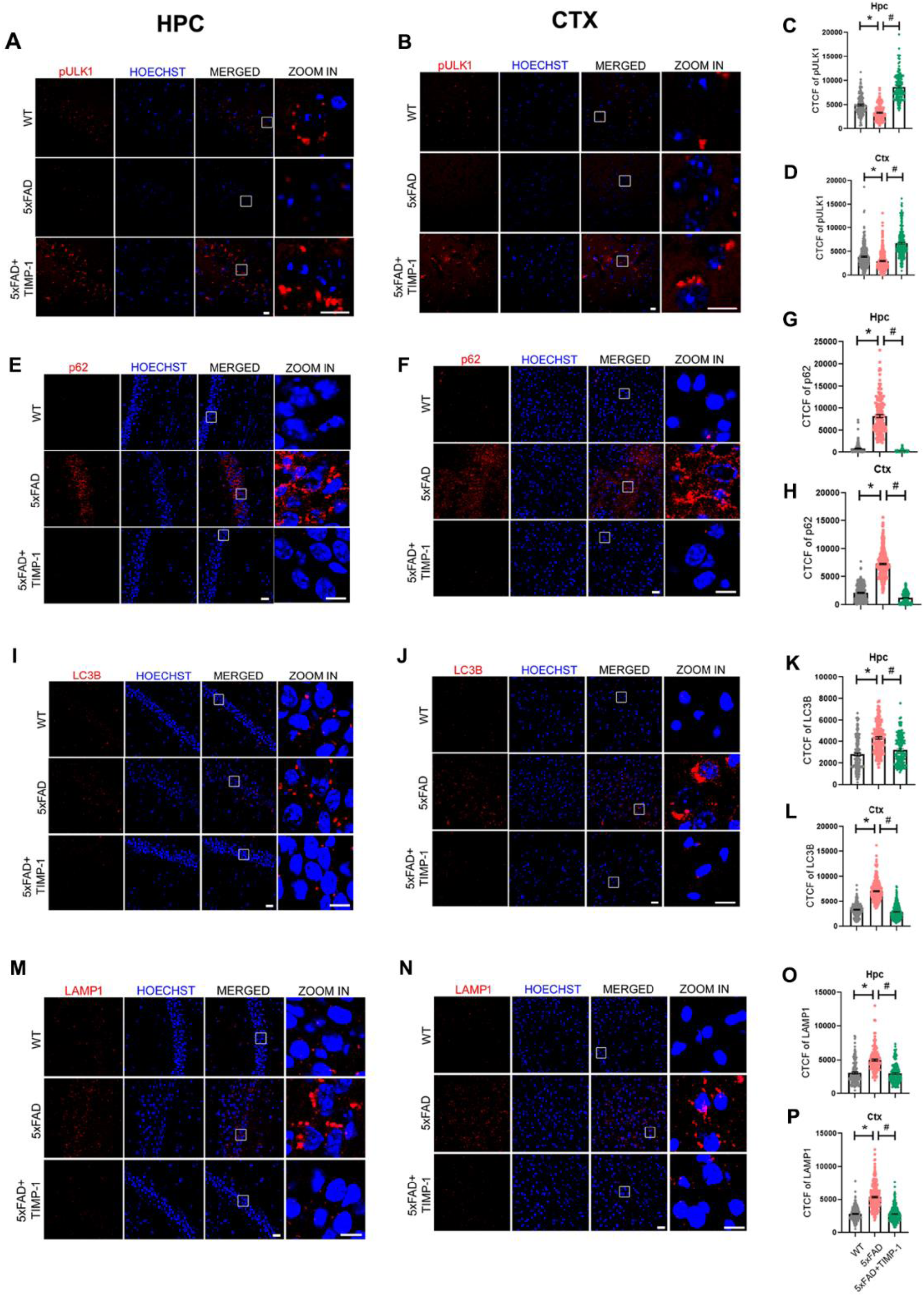
TIMP-1 injection in 5xFAD mice protects against impaired autophagy. **A-B, E-F, I-J, M-N,** Representative immunofluorescence images for pULK1(S757) (**A**, Hpc; **B**, Ctx), p62 (**E**, Hpc; **F**, Ctx), LC3B (**I**, Hpc; **J**, Ctx) and LAMP1 (**M**, Hpc; **N**, Ctx), 14 days post TIMP-1 injection in 5xFAD mice. White boxes in MERGED panel demarcates the region-of-interest for higher magnification images shown in the right panel. **C-D, G-H, K-L, O-P**, CTCF for pULK1(S757) (**C**, Hpc; **D,** Ctx), p62 (**G**, Hpc; **H,** Ctx), LC3B (**K,** Hpc**; L,** Ctx) and LAMP1 (**O,** Hpc**; P,** Ctx). N=3 mice/group, n=50 cells/mouse from hippocampus (Hpc) and 100 cells/mouse from cortex (Ctx) quantified from 40x images, One-way ANOVA, Tukey’s post hoc, mean±s.e.m.*P<0.0001, ^#^P<0.0001; Scale bar=25 µm.

**Figure S6:**
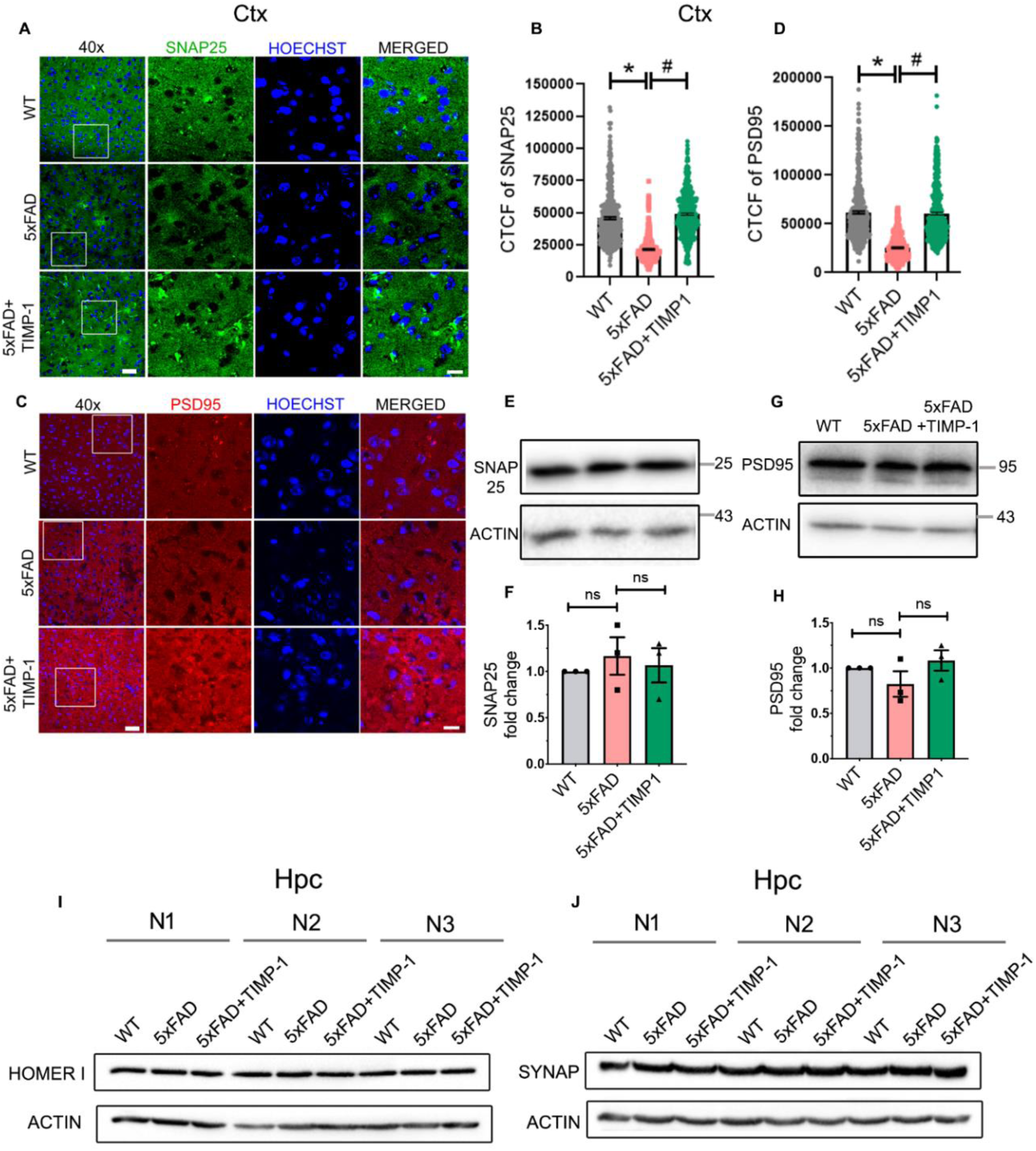
TIMP-1 differentially affects synaptic protein expressions in cortex and hippocampus of 5xFAD mice. **A,C,** Representative immunofluorescence images for SNAP25 (green, **A**) and PSD95 (red, **C**) in coronal sections of WT, 5xFAD and 5xFAD+TIMP-1 cortex; scale bar, 30 µm at 40x magnification (white box indicates the region selected for further zoom-in). Zoom-in merged images are shown with nuclei stained with Hoechst (blue); scale bar, 15µm. **B,D**, CTCF calculated forfor SNAP25 (**B**) and PSD95 (**D**), N=3 animals/group, n=150 cells/animal from Cortex (Ctx) quantified from 40x images; One-way ANOVA, Tukey’s post hoc, mean±s.e.m. *P<0.0001, **^#^**P<0.0001 for **B** and **D**. Representative immunoblots for SNAP25 (**E**) and PSD95 (**G**) from Ctx whole tissue lysates collected 14 days post TIMP-1 injection. Densitometric analyses and fold changes for normalized SNAP25 (**F**) and PSD95 (**H**) w.r.t. to WT group. N = 3 animals/group; One-way ANOVA, Tukey’s post hoc, mean±s.e.m. ns=not significant. **I-J**, Representative immunoblots for Homer-1 (**I**) and Synaptophysin-1 (SYNAP, **J**) from whole hippocampal (Hpc) tissue of mice 14 days post TIMP-1 injection. N=animal number.

**Figure S7:**
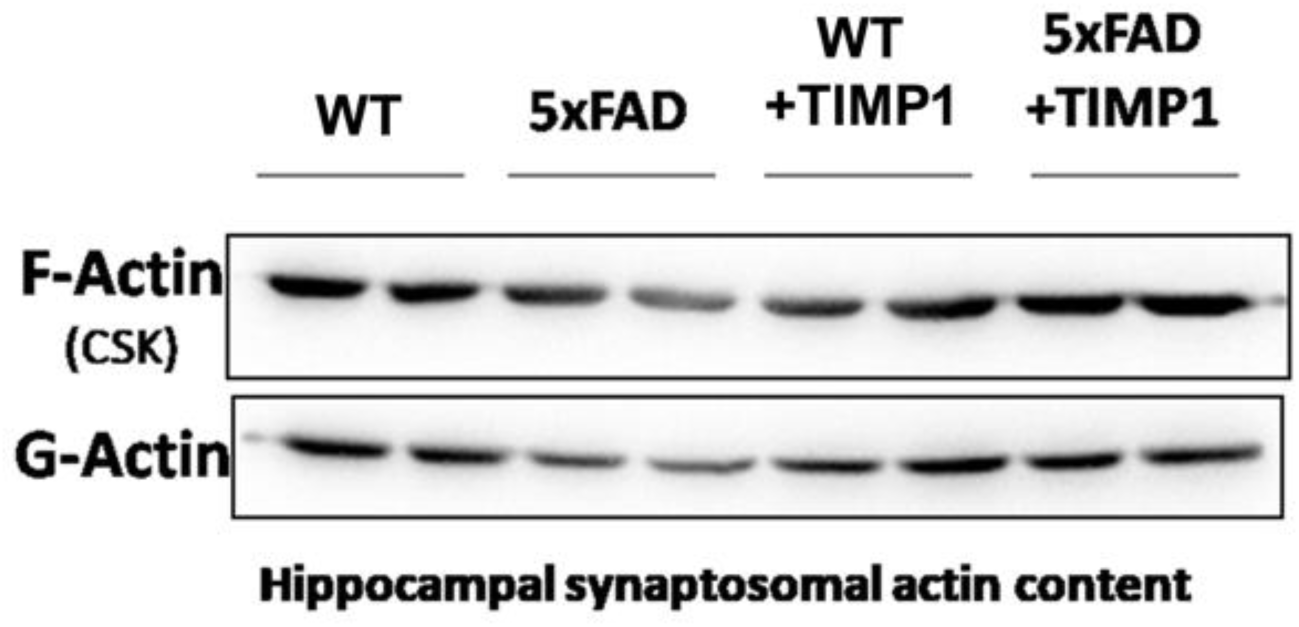
TIMP-1 injection does not alter F/G actin ratio in WT mice. Representative immunoblots for F-Actin and G-Actin from hippocampal synaptosomes isolated from the four treatment groups, 14 days post TIMP-1 injection. CSK=cytoskeletal fraction of synaptosome.

**Figure S8:**
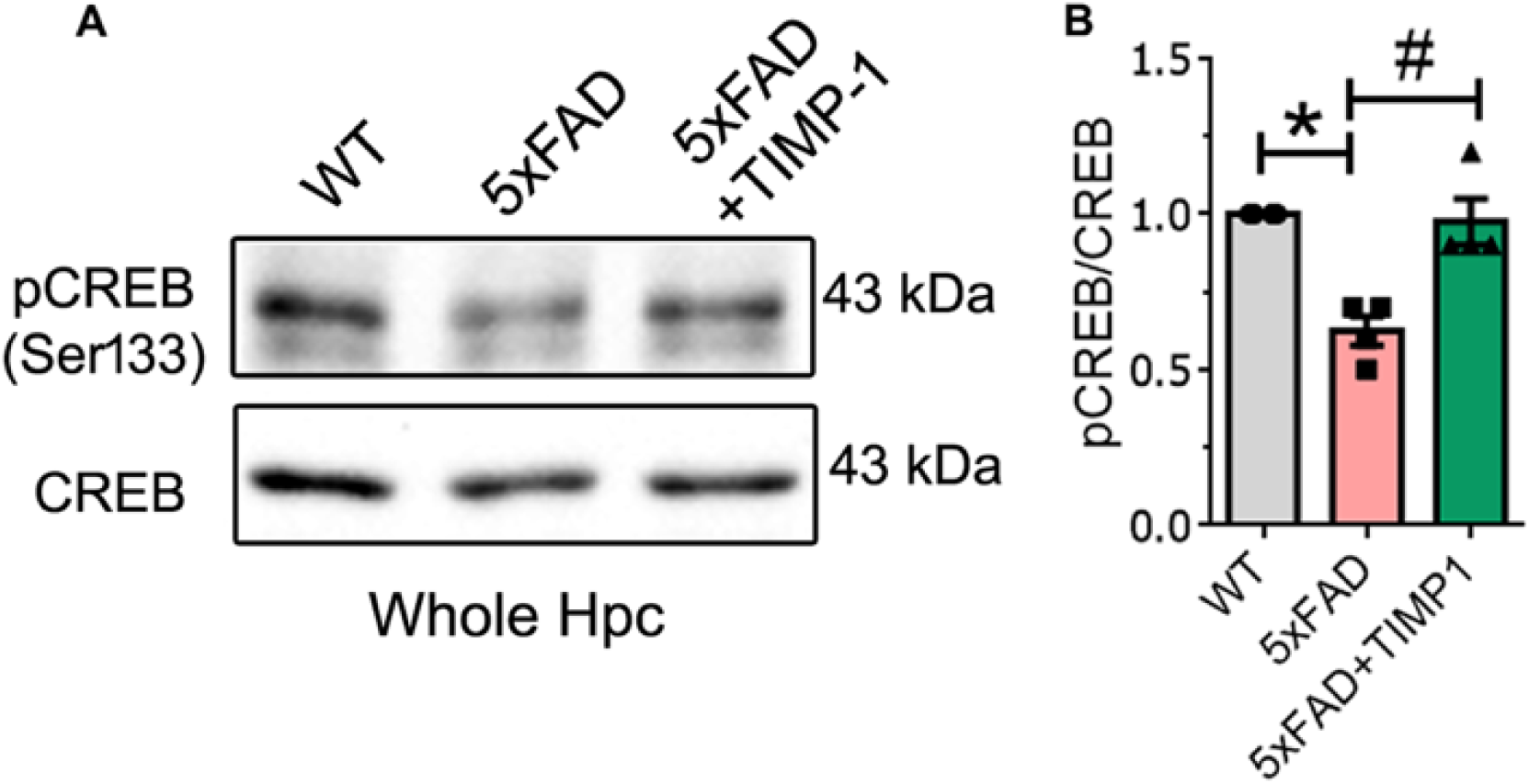
TIMP-1 improves CREB activity in the hippocampus of 5xFAD mice. **A,** Representative immunoblots for pCREB(Ser133) and CREB from hippocampal whole tissue isolated 14 days post TIMP-1 treatment. **B**, Densitometric analyses for pCREB/CREB showing fold change w.r.t. WT group. N=4 mice/group; One-way ANOVA, Tukey’s post hoc, mean±s.e.m.; *P<0.0015, **^#^**P<0.0024.

**Figure S9:**
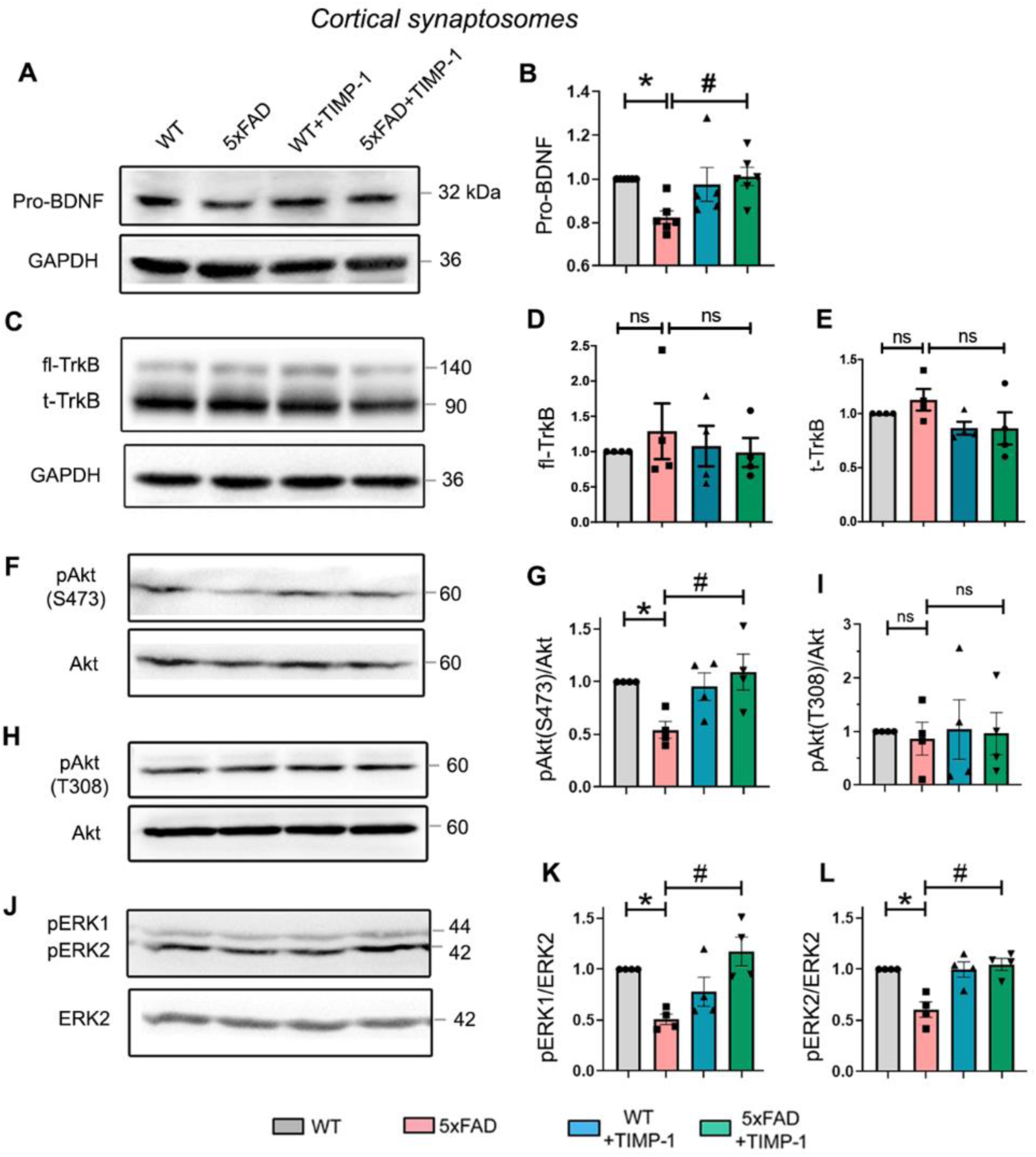
TIMP-1 improves BDNF and related signaling at cortical synapses of 5xFAD mice. WB data from cortical synaptosomes 14 days post TIMP-1 treatment. Representative blots for Pro-BDNF (**A**), TrkB (**C**), pAkt(S473) (**F**), pAkt(T308) (**H**) and pERK1/2 (**J**) with corresponding loading controls. Densitometric analyses and normalized fold change w.r.t. WT for Pro-BDNF (**B**), fl-TrkB (**D**), t-TrkB (**E**), pAkt(S473)/Akt (**G**), pAkt(t308)/Akt (**I**), pERK1/ERK2 (**K**) and pERK2/ERK2 (**L**). N=6 independent synaptosome isolations/group, One-way ANOVA, Tukey’s post hoc for **B**; N=4 independent synaptosome isolations/group, One-way ANOVA, Tukey’s post hoc for **D,E,G,I,K,L**. Each synaptosome batch (N) isolated from 6-7 animals for each group. All data expressed as mean±s.e.m. Fl-TrkB, Full-length TrkB; t-TrkB, truncated TrkB. *P<0.05 versus WT, ^#^P<0.05 versus 5xFAD, ns=not significant. Exact P values for this figure are given in Table S1.

### Supplementary Table

**Table S1: Overview statistics for Figures 1, 5, 7, S2 and S9**.

