## Supplementary Table 1 for "An astrocyte-derived cytokine TIMP-1 restores synaptic plasticity in an Alzheimer’s disease model"

| Table S1. Overview statistics for Figures 1, 5, 7, S2 and S9. | | | |
|---|---|---|---|
| **Figure** | **Data Type** | **Sample size (N, n) and P value** | **Statistical test** |
| 1B | Mean ± s.e.m. | N = 6 animals/group/age | Two-tailed unpaired t-test |
| | | 2M: WT versus 5xFAD, P=0.0333 | |
| | | 4M: WT versus 5xFAD, P<0.0001 | |
| | | 6M: WT versus 5xFAD, P<0.0001 | |
| | | 8M: WT versus 5xFAD, P<0.0001 | |
| 1E | Mean ± s.e.m. | N = 3 independent experiments, n = 3 slices/animal, P<0.0001 | Two-tailed unpaired t-test |
| 1G-H, S2A-B | | WT: N = 16 animals | |
| | | 5xFAD: N = 10 animals | |
| | | 5xFAD+aCSF: N = 12 animals | |
| | | 5xFAD+TIMP-1: N = 10 animals | |
| 1G | Mean ± s.e.m. | WT versus 5xFAD, P=0.0001 | One-way ANOVA followed by Tukey's post hoc test |
| | | WT versus 5xFAD+aCSF, P=0.0002 | |
| | | 5xFAD versus 5xFAD+TIMP-1, P=0.0458 | |
| | | 5xFAD+aCSF versus 5xFAD+TIMP-1, P=0.0771 | |
| 1H | Mean ± s.e.m. | WT versus 5xFAD, P<0.0001 | One-way ANOVA followed by Tukey's post hoc test |
| | | WT versus 5xFAD+aCSF, P=0.0009 | |
| | | 5xFAD versus 5xFAD+TIMP-1, P<0.0001 | |
| | | 5xFAD+aCSF versus 5xFAD+TIMP-1, P=0.0005 | |
| S2A | Mean ± s.e.m. | WT versus 5xFAD, P<0.0001 | One-way ANOVA followed by Tukey's post hoc test |
| | | WT versus 5xFAD+aCSF, P<0.0001 | |
| | | 5xFAD versus 5xFAD+TIMP-1, P=0.0047 | |
| | | 5xFAD+aCSF versus 5xFAD+TIMP-1, P<0.0001 | |
| S2B | Mean ± s.e.m. | WT versus 5xFAD, P<0.0001 | One-way ANOVA followed by Tukey's post hoc test |
| | | WT versus 5xFAD+aCSF, P=0.0008 | |
| | | 5xFAD versus 5xFAD+TIMP-1, P<0.0001 | |
| | | 5xFAD+aCSF versus 5xFAD+TIMP-1, P=0.0005 | |
| 1J, S2C-D | Mean ± s.e.m. | WT: N = 18 animals | |
| | | 5xFAD: N = 14 animals | |
| | | 5xFAD+aCSF: N = 10 animals | |
| | | 5xFAD+TIMP-1: N = 14 animals | |
| | | WT(ITL) versus WT (RTL), P<0.0001 | Two-tailed unpaired t-test |
| | | 5xFAD (ITL) versus 5xFAD (RTL), P=0.1650 | |
| | | 5xFAD+aCSF (ITL) versus 5xFAD+aCSF (RTL), P=0.8062 | |
| | | 5xFAD+TIMP-1 (ITL) versus 5xFAD+TIMP-1 (RTL), P<0.0001 | |
| | | WT (ITL-RTL) versus 5xFAD (ITL-RTL), P<0.0001 | One-way ANOVA |
| | | WT (ITL-RTL) versus 5xFAD+aCSF (ITL-RTL), P=0.0356 | |

| | | | |
|---|---|---|---|
| | | 5xFAD (ITL-RTL) versus 5xFAD+TIMP-1 (ITL-RTL), P<0.0001 | followed by Tukey's post hoc test |
| | | 5xFAD+aCSF (ITL-RTL) versus 5xFAD+TIMP-1 (ITL-RTL), P=0.0026 | |
| 1L-N, S2E | Mean ± s.e.m. | WT: N = 19 animals | |
| | | 5xFAD: N = 12 animals | |
| | | 5xFAD+aCSF: N = 11 animals | |
| | | 5xFAD+TIMP-1: N = 14 animals | |
| 1L, S2E | | WT(F) versus WT (N), P=0.0006 | Two-tailed unpaired t-test |
| | | 5xFAD (F) versus 5xFAD (N), P=0.4145 | |
| | | 5xFAD+aCSF (F) versus 5xFAD+aCSF (N), P=0.4667 | |
| | | 5xFAD+TIMP-1 (F) versus 5xFAD+TIMP-1 (N), P=0.0018 | |
| 1M | | WT versus 5xFAD, P<0.0001 | One-way ANOVA followed by Tukey's post hoc test |
| | | WT versus 5xFAD+aCSF, P<0.0001 | |
| | | 5xFAD versus 5xFAD+TIMP-1, P=0.0008 | |
| | | 5xFAD+aCSF versus 5xFAD+TIMP-1, P=0.0003 | |
| 1N | | WT versus 5xFAD, P<0.0001 | One-way ANOVA followed by Tukey's post hoc test |
| | | WT versus 5xFAD+aCSF, P<0.0001 | |
| | | 5xFAD versus 5xFAD+TIMP-1, P=0.0008 | |
| | | 5xFAD+aCSF versus 5xFAD+TIMP-1, P=0.0003 | |
| 1P | Mean ± s.e.m. | WT: N = 14 animals | One-way ANOVA followed by Tukey's post hoc test |
| | | 5xFAD: N = 12 animals | |
| | | 5xFAD+aCSF: N = 8 animals | |
| | | 5xFAD+TIMP-1: N = 12 animals | |
| | | WT versus 5xFAD, P=0.0098 | |
| | | WT versus 5xFAD+aCSF, P=0.0068 | |
| | | 5xFAD versus 5xFAD+TIMP-1, P<0.0001 | |
| | | 5xFAD+aCSF versus 5xFAD+TIMP-1, P<0.0001 | |
| 1R | Mean ± s.e.m. | WT: N = 18 animals | One-way ANOVA followed by Tukey's post hoc test |
| | | 5xFAD: N = 14 animals | |
| | | 5xFAD+aCSF: N = 8 animals | |
| | | 5xFAD+TIMP-1: N = 12 animals | |
| | | WT versus 5xFAD, P=0.0009 | |
| | | WT versus 5xFAD+aCSF, P=0.0024 | |
| | | 5xFAD versus 5xFAD+TIMP-1, P<0.0001 | |
| | | 5xFAD+aCSF versus 5xFAD+TIMP-1, P<0.0001 | |
| 5K | Mean ± s.e.m. | SNAP25: Ctx, N = 4 independent synaptosome isolations | One-way ANOVA followed by Tukey's post hoc test |
| | | WT versus 5xFAD, P=0.0012 | |
| | | 5xFAD versus 5xFAD+TIMP-1, P=0.0001 | |
| | | SNAP25: Hpc, N = 4 independent synaptosome isolations | One-way ANOVA |

| | | | |
|---|---|---|---|
| | | WT versus 5xFAD, P=0.0049 | followed by Tukey's post hoc test |
| | | 5xFAD versus 5xFAD+TIMP-1, P=0.0154 | |
| 5L | Mean ± s.e.m. | SYNAP: Ctx, N = 3 independent synaptosome isolations | Two-tailed unpaired t-test |
| | | WT versus 5xFAD, P=0.0354 | |
| | | 5xFAD versus 5xFAD+TIMP-1, P=0.2050 | |
| | | SYNAP: Hpc, N = 4 independent synaptosome isolations | Two-tailed unpaired t-test |
| | | WT versus 5xFAD, P=0.0103 | |
| | | 5xFAD versus 5xFAD+TIMP-1, P=0.1229 | |
| | | One batch among these synaptosomal isolations (for each region) is devoid of WT+TIMP-1 group | |
| 5M | Mean ± s.e.m. | PSD95: Ctx, N = 5 independent synaptosome isolations | Two-tailed unpaired t-test |
| | | WT versus 5xFAD, P=0.0007 | |
| | | 5xFAD versus 5xFAD+TIMP-1, P=0.0397 | |
| | | PSD95: Hpc, N = 4 independent synaptosome isolations | Two-tailed unpaired t-test |
| | | WT versus 5xFAD, P=0.0008 | |
| | | 5xFAD versus 5xFAD+TIMP-1, P=0.0260 | |
| | | One batch among Ctx synaptosomal isolations is devoid of WT+TIMP-1 group | |
| 5N | Mean ± s.e.m. | HOMER I: Ctx, N = 4 independent synaptosome isolations | Two-tailed unpaired t-test |
| | | WT versus 5xFAD, P=0.0003 | |
| | | 5xFAD versus 5xFAD+TIMP-1, P=0.0074 | |
| | | HOMER I: Hpc, N = 3 independent synaptosome isolations | Two-tailed unpaired t-test |
| | | WT versus 5xFAD, P=0.0053 | |
| | | 5xFAD versus 5xFAD+TIMP-1, P=0.1969 | |
| 7B | Mean ± s.e.m. | Pro-BDNF: N = 4 independent experiments | One-way ANOVA followed by Tukey's post hoc test |
| | | WT versus 5xFAD, P=0.0157 | |
| | | 5xFAD versus 5xFAD+TIMP-1, P=0.0189 | |
| 7D | Mean ± s.e.m. | fl-TrkB: N = 3 independent experiments | One-way ANOVA followed by Tukey's post hoc test |
| | | WT versus 5xFAD, P=0.2410 | |
| | | 5xFAD versus 5xFAD+TIMP-1, P=0.0769 | |
| 7E | Mean ± s.e.m. | t-TrkB: N = 3 independent experiments | One-way ANOVA followed by Tukey's post hoc test |
| | | WT versus 5xFAD, P=0.9811 | |
| | | 5xFAD versus 5xFAD+TIMP-1, P=0.7921 | |

| | | | |
|---|---|---|---|
| 7G | Mean ± s.e.m. | pAkt(S473): N = 4 independent experiments<br>WT versus 5xFAD, P=0.0388<br>5xFAD versus 5xFAD+TIMP-1, P=0.0042 | One-way ANOVA followed by Tukey's post hoc test |
| 7I | Mean ± s.e.m. | pAkt(T308): N = 4 independent experiments<br>WT versus 5xFAD, P=0.0464<br>5xFAD versus 5xFAD+TIMP-1, P=0.0108 | One-way ANOVA followed by Tukey's post hoc test |
| 7K | Mean ± s.e.m. | pERK1: N = 4 independent experiments<br>WT versus 5xFAD, P=0.3004<br>5xFAD versus 5xFAD+TIMP-1, P=0.3004 | One-way ANOVA followed by Tukey's post hoc test |
| 7L | Mean ± s.e.m. | pERK2: N = 4 independent experiments<br>WT versus 5xFAD, P=0.9880<br>5xFAD versus 5xFAD+TIMP-1, P=0.8282 | One-way ANOVA followed by Tukey's post hoc test |
| 7N | Mean ± s.e.m. | Pro-BDNF: N = 7 independent synaptosome isolations<br>WT versus 5xFAD, P=0.0269<br>5xFAD versus 5xFAD+TIMP-1, P=0.0076<br>One batch among synaptosomal isolations is devoid of WT+TIMP-1 group | Mixed effect ANOVA analysis followed by Tukey's post hoc test |
| 7P | Mean ± s.e.m. | fl-TrkB: N = 4 independent synaptosome isolations<br>WT versus 5xFAD, P=0.8322<br>5xFAD versus 5xFAD+TIMP-1, P=0.3841 | One-way ANOVA followed by Tukey's post hoc test |
| 7Q | Mean ± s.e.m. | t-TrkB: N = 4 independent synaptosome isolations<br>WT versus 5xFAD, P=0.9742<br>5xFAD versus 5xFAD+TIMP-1, P=0.9210 | One-way ANOVA followed by Tukey's post hoc test |
| 7S | Mean ± s.e.m. | pAkt(S473): N = 5 independent synaptosome isolations<br>WT versus 5xFAD, P=0.0023<br>5xFAD versus 5xFAD+TIMP-1, P=0.0346 | One-way ANOVA followed by Tukey's post hoc test |
| 7U | Mean ± s.e.m. | pAkt(T308): N = 4 independent synaptosome isolations<br>WT versus 5xFAD, P=0.0262<br>5xFAD versus 5xFAD+TIMP-1, P=0.3264 | One-way ANOVA followed by Tukey's post hoc test |
| 7W | | pERK1: N = 6 independent synaptosome isolations | |

| | | | |
|---|---|---|---|
| | Mean ± s.e.m. | WT versus 5xFAD, P=0.0285<br>5xFAD versus 5xFAD+TIMP-1, P=0.0222 | One-way ANOVA followed by Tukey's post hoc test |
| 7X | Mean ± s.e.m. | pERK2: N = 6 independent synaptosome isolations<br>WT versus 5xFAD, P=0.0469<br>5xFAD versus 5xFAD+TIMP-1, P=0.0405 | One-way ANOVA followed by Tukey's post hoc test |
| S9B | Mean ± s.e.m. | Pro-BDNF: N = 6 independent synaptosome isolations<br>WT versus 5xFAD, P=0.0358<br>5xFAD versus 5xFAD+TIMP-1, P=0.0244<br>One batch among synaptosomal isolations is devoid of WT+TIMP-1 group | One-way ANOVA followed by Tukey's post hoc test |
| S9D | Mean ± s.e.m. | fl-TrkB: N = 4 independent synaptosome isolations<br>WT versus 5xFAD, P=0.8672<br>5xFAD versus 5xFAD+TIMP-1, P=0.8524 | One-way ANOVA followed by Tukey's post hoc test |
| S9E | Mean ± s.e.m. | t-TrkB: N = 4 independent synaptosome isolations<br>WT versus 5xFAD, P=0.7758<br>5xFAD versus 5xFAD+TIMP-1, P=0.2454 | One-way ANOVA followed by Tukey's post hoc test |
| S9G | Mean ± s.e.m. | pAkt(S473): N = 4 independent synaptosome isolations<br>WT versus 5xFAD, P=0.0371<br>5xFAD versus 5xFAD+TIMP-1, P=0.0146<br>One batch among synaptosomal isolations is devoid of WT+TIMP-1 group | One-way ANOVA followed by Tukey's post hoc test |
| S9I | Mean ± s.e.m. | pAkt(T308): N = 4 independent synaptosome isolations<br>WT versus 5xFAD, P=0.9942<br>5xFAD versus 5xFAD+TIMP-1, P=0.9980 | One-way ANOVA followed by Tukey's post hoc test |
| S9K | Mean ± s.e.m. | pERK1: N = 4 independent synaptosome isolations<br>WT versus 5xFAD, P=0.0258<br>5xFAD versus 5xFAD+TIMP-1, P=0.0033 | One-way ANOVA followed by Tukey's post hoc test |
| S9L | Mean ± s.e.m. | pERK2: N = 4 independent synaptosome isolations<br>WT versus 5xFAD, P=0.0030<br>5xFAD versus 5xFAD+TIMP-1, P=0.0012 | One-way ANOVA followed by Tukey's post hoc test |
